# Experimental evolution of *Escherichia coli* on semi-dry silver, copper, stainless steel, and glass surfaces

**DOI:** 10.1101/2024.06.06.597739

**Authors:** Merilin Rosenberg, Sandra Park, Sigrit Umerov, Angela Ivask

**Affiliations:** Institute of Molecular and Cell Biology, University of Tartu, Riia 23, 51010 Tartu, Estoni

## Abstract

To study bacterial adaptation to antimicrobial metal surfaces in application-relevant conditions, *Escherichia coli* was exposed to copper and silver surfaces for thirty exposure cycles in low-organic dry or high-organic humid conditions. The evolved populations demonstrated increased metal surface tolerance without concurrent increase in MBC and MIC values of respective metal ions or selected antibiotics. Mutation analysis did not detect increased mutation accumulation nor mutations in *cop, cus, cue, sil, pco* or general efflux genes known to actively maintain copper/silver homeostasis. Instead, during cyclic exposure mutations in genes related to cellular barrier functions and sulfur metabolism were enriched potentially suggesting that reducing bioavailability and passively restricting uptake of the toxic metals rather than active efflux is selected for on copper and silver surfaces. The changes detected in the evolved populations did not indicate an increased risk of antibiotic cross-resistance as a result of copper or silver surface exposure. However, rapid emergence of mutations in *silS* activated the cryptic *sil* efflux locus during silver ion challenge in liquid MBC assay with the evolved populations. The *silS* mutants showed no benefit on copper and silver surfaces but demonstrated decreased sensitivity to ampicillin and ciprofloxacin as well as copper and silver ions in liquid tests indicating that efflux might be specific to granting heavy metal tolerance in liquid but not surface exposure format. Our findings highlight the critical importance of appropriate exposure conditions not only in efficacy assessment but also risk assessment of antimicrobial surface applications.

**Importance:** This study examines the evolutionary adaptations of *Escherichia coli* after semi-dry exposure to copper and silver surfaces, leading to an increase in surface tolerance but no increase in mutation accumulation or substantially enhanced metal ion tolerance in standard tests. Notably, enriched mutations indicate a shift toward more energy-passive mechanisms of metal tolerance. Additionally, while enhanced silver efflux was rapidly selected for in a single round of silver exposure in liquid tests and substantially increased copper and silver ion tolerance in conventional test formats, the causal mutations did not improve viability on silver and copper surfaces, underscoring the different fitness scenarios of tolerance mechanisms dependent on exposure conditions. These findings emphasize the need for appropriate exposure conditions in evaluating of both efficacy and the potential risks of using antimicrobial surfaces, as the results from conventional liquid-based tests may not apply in solid contexts.

## Introduction

Antimicrobial surfaces are expected to passively reduce bioburden on surfaces with elevated risk of pathogen transmission. Among the materials used, silver and copper based applications have the longest application history as well as wide commercial use [1]. Despite challenges in efficacy assessment of such materials in environmentally exposed conditions there is a wide range of environmentally directed efficacy scenarios with highly variable proportions of microbial survivors from quick killing of most microbes to effectively no biocidal activity [2]. As with any antimicrobial application, the possibility of high numbers of survivors has raised a growing concern that prolonged or repeated exposure to antimicrobial surfaces could not only increase biocide tolerance but also enhance development of antibiotic resistance in these microbes.

Increased risk of antibiotic resistance could arise from co-selection, cross-selection or co-regulation of molecular mechanisms that increase both antimicrobial metal as well as antibiotic tolerance while increased mutation rates could lead to quicker adaptations leading to the development of these mechanisms. The risk of co-selection of metal tolerance and antibiotic resistance is obvious and inevitable in the case of large multiresistance plasmids [3], [4] but in case of cross-resistance and co-regulation, test conditions might substantially affect selection. Mechanisms of ionic silver and copper tolerance partly overlap and are mainly associated with energy-expensive active ion efflux systems in case of silver (*e.g.* the Sil, Cus, Cop, Pco systems) [5] whereas employment of additional defense mechanisms is needed in case of the redox-active copper [6]. Some of the potential tolerance mechanisms, especially enhanced efflux systems, can also lead to antibiotic cross-resistance. Although cross-selection of mutations in efflux systems and outer membrane proteins has been well described in decreased sensitivity to metals and antibiotics (*e.g.* [7], [8], [9]) the risk of *de novo* resistance development, potential cross-selection of beneficial traits in multistress exposure, and effect of non-genetic co-regulation are more challenging to study.

Mutation rate of an evolving population can naturally fluctuate in time due to the fixation of various hypermutator and antimutator alleles [10], [11], [12] [13]. At the same time, presence of the hypermutator phenotype has been associated with increased pathogenicity [14], [15], [16], [17], [18], [19] and antibiotic resistance development [20], [21], [22], [23]. As shown previously, in growth-supporting exposure media subinhibitory copper and silver could increase mutation rate [24], [25] and heavy metals could thereby lead to antibiotic resistance development [26]. However, the role of metal-based biocides in *de novo* antibiotic resistance development or their effect on mutation rate is controversial, especially in lethal and/or low-organic conditions. Similarly to antimicrobial surface efficacy, molecular defense mechanisms potentially leading to cross-resistance or co-regulation are possibly highly dependent on biotic and abiotic environment as well as the resources available to the bacteria. While it has been feared that antimicrobial agents that increase mutation frequency could collaterally enhance antibiotic resistance development and increase pathogenicity there are also examples where the contrary is demonstrated and some metal ions such as copper and zinc can suppress not only hypermutation but also conjugative gene transfer possibly by blocking the SOS response [27], [28], [29]. In addition, one must keep in mind that the majority of the published data is acquired from liquid test formats which do not necessarily resemble surface properties other than release of the active ingredient. Instead, the antimicrobial effect on *e.g.*, copper-based surfaces is rather dependent on physical properties of the surface material such as its copper content of the alloy and direct surface-microbe contact not the amount copper released from the surface [30]. It has also been shown that the speed of contact-killing on copper surfaces is linked to bacterial cell surface components and their interactions with the metal surface indicating limited ability to evolve resistance to metallic copper [31]. On air-exposed solid surfaces, bacteria are not only exposed to the biocide incorporated into the surface material but also several other environmental stressors such as desiccation, light exposure, osmotic shock, starvation *etc*. The specific stress responses mechanistically overlap with response to metal ions (Supplementary Fig S1) highlighting possible co-regulational toll or benefit associated with combined exposure. In contrast to subinhibitory copper concentrations in liquid growth medium, transient exposure to copper surfaces in environmentally exposed non-growth supporting conditions has been shown not to increase mutation frequency of *E. coli* [32].

The evolution of antimicrobial tolerance on solid surfaces is a rather novel topic with two articles recently published on the matter. The two studies by Bleichert *et al*. [33] and Xu *et al*. [34] both exposed bacteria to copper-based surfaces in semi-dry conditions. However, they investigated evolution of different bacterial species on copper-based surfaces using different approaches to controls, exposure and growth conditions which complicates comparing the results that largely differ in mutation accumulation as well as changes in copper tolerance. Xu *et al*. used passaging *Pseudomonas fluorescens* in liquid medium as a negative control for brass and copper surface exposure and used liquid culture for survivor cultivation between exposures thereby lacking a control for exposure conditions and causing competition for growth rate between exposures. They analyzed ≥50% fixed single nucleotide variations (SNVs) and found a large number of mutations (incl. in *cop* and *cue* genes) as well as increased mutation accumulation on copper surfaces compared to negative control. Bleichert *et al*. used stainless steel surface exposure as a negative control for *Escherichia coli* and *Staphylococcus aureus* and propagated survivors on solid medium between exposures but also introduced an additional bottleneck by selecting only some survivor colonies to start the next exposure cycle. They mainly analyzed ≥ 80% fixed SNVs, found very few mutations and did not identify mutations explaining changes in copper tolerance of the evolved bacteria.

The rather inconsistent results in the literature on the effects of metal exposure on metal tolerance, antibiotic cross-resistance and bacterial mutation rates reveal a knowledge gap in how exposure conditions affect bacterial adaptation to antimicrobial metal ions versus respective solid metal surfaces in more realistic semi-dry exposure conditions. With the current study we aimed to evaluate bacterial adaptation as well as associated risks of potential development of antibiotic cross-resistance in response to repeated exposure of *Escherichia coli* to silver and copper surfaces in different application-relevant semi-dry exposure conditions.

## Materials and Methods

### Strains and media

*Escherichia coli* strain routinely used for efficacy testing of antibacterial agents was used in all experiments. *E. coli* DSM 1576 (ATCC 8739, NCIB 8545, WDCM 00012, Crooks strain) was freshly ordered from the DSMZ (German Collection of Microorganisms and Cell Cultures GmbH). 2^nd^ subculture of the received strain on LB agar medium (Table 1) was used for the evolution experiments. Here onwards, the more widely used synonymous ATCC 8739 strain number is used.

**Table 1.** Experimental conditions used in the evolution experiments.

|  | Low-organic dry (LD) conditions |  |  |  | High-organic humid (HH) conditions |  |  |  |
| --- | --- | --- | --- | --- | --- | --- | --- | --- |
| Organism | Escherichia coli ATCC 8739 |  |  |  |  |  |  |  |
| Inoculum cell count | 8.1x10<sup>6</sup> CFU per surface |  |  |  |  |  |  |  |
| Exposure temperature | 22°C |  |  |  |  |  |  |  |
| Exposure medium | 500-fold diluted nutrient broth [37] (NB: 0.026 g/l organics):<br>0.006 g/l meat extract<br>0.02 g/l peptone<br>0.01 g/l NaCl |  |  |  | Organic proportion of EPA soil load [36](SL: 6.8 g/l organics):<br>bovine serum albumin 2.5 g/l<br>yeast extract 3.5 g/l<br>mucin 0.8 g/l |  |  |  |
| Inoculum volume | 5 x 2 µl |  |  |  | 1 x 50 µl |  |  |  |
| Relative humidity | 50% |  |  |  | 90% |  |  |  |
| Surface material | Copper | Silver | Steel | Glass | Copper | Silver | Steel | Glass |
| Exposure duration | 20-25 min | 2 h | 2 h | 2 h | 1 h | 6 h | 6 h | 6 h |
| Toxicity-neutralizing harvesting medium | Soy casein digest lecithin polysorbate broth (SCDLP [37]):<br>17 g/l casein peptone, 3 g/l soybean peptone, 5 g/l NaCl, 2.5 g/l Na<sub>2</sub>HPO<sub>4</sub>, 2.5 g/l glucose, 1.0 g/l lecithin, 7.0 g/l Tween80 |  |  |  |  |  |  |  |
| Medium for clonal growth between exposures | LB agar medium: 5 g/l yeast extract, 10 g/l tryptone, 5 g/l NaCl, 15 g/l agar |  |  |  |  |  |  |  |
| Time of clonal growth between exposures | ~45 h |  |  |  | ~40 h |  |  |  |
| Harvesting medium for populations from clonal growth plates for reinoculation | Phosphate-buffered saline (PBS):<br>10-fold diluted stock of: 8 g/l NaCl, 0.2 g/l KCl, 1.44 g/l Na<sub>2</sub>HPO<sub>4</sub>, 0.2 g/l KH<sub>2</sub>PO<sub>4</sub>; pH 7.1 |  |  |  |  |  |  |  |

### Surface treatment

99% copper (Metroprint OY, Estonia) and 99.95% silver (Surepure Chemetals, USA) were used as historically well-known antimicrobial materials. Stainless steel (AISI 304 SS, incl. 18% Cr, 8% Ni, 1.4% Mn; 2B finish; Aperam-Stainless, France) is a widely used metallic surface and has been frequently suggested as a control surface in studies involving metal alloys as well as standard biocide testing methods. 18x18 mm borosilicate glass microscopy cover slips were used as a biologically inert control surface (Corning Inc., USA). Steel, copper, and silver metal sheets were laser-cut to Ø 2 cm coupons and reused in evolution experiments. After each exposure the metal coupons were collected by type and inactivated in 70% ethanol for 15 minutes. The cleaning cycle consisted of 3 sequential solution steps (5% citric acid, acetone, 70% EtOH) with glass beads and 2 rinsing steps without glass beads. Solutions steps included ultrasonication in water bath for 15 minutes followed by shaking for 2 minutes manually before and after ultrasonication. Between each solution step the coupons were rinsed in distilled water by shaking. After cleaning the coupons were dried in a biosafety cabinet and stored in sterile Petri dishes. Glass surfaces were rinsed with 70% EtOH, dried and exposed to UV-C radiation for 15 min on both sides.

### Evolution experiments

The evolution experiments in low-organic dry (LD) and high-organic humid (HH) conditions (Table 1) each consisted of three parts: (re-)inoculation, exposure in the climate chamber and harvesting of cells for plating and reinoculation repeated in 30 cycles. In each cycle, a sample from each inoculum was preserved in glycerol stock at -80°C to prevent loss of lineages due to contamination and/or lethal exposure. After the 30^th^ cycle the collected colony suspension was also stored at –20° C for DNA extraction. The two evolution experiments were started from single colonies of *E. coli* ATCC 8739 and cycled 5 parallel populations on each surface type resulting in 40 evolved populations.

For cyclic exposure sterile test surfaces were placed into the wells of a 6-well plate. In case of humid exposure, a sterile 3D printed plastic grid was used to suspend each test surface above 3 ml of sterile water pipetted below the suspension grid to preserve moisture in the system.

For the first cycle of each experiment *E. coli* ATCC 8739 was inoculated on LB agar plates from glycerol stock culture and incubated for 24 h at 37° C from which a single colony was further inoculated on LB agar for 20 h 37° C. From the resulting second subculture all colonies were collected by suspending with 5 ml of phosphate buffered saline (PBS). 500 µl of that suspension was preserved in glycerol stock and sequenced as the “ancestor”. For 2^nd^ to 30^th^ exposure cycle, inoculum was made from resuspended colonies from the previous cycle plate-out. Prior to inoculation, the cell density was spectrophotometrically adjusted to 3.3x10^8^ CFU/ml or 1.6x10^9^ CFU/ml [35] for exposure with EPA soil load (SL, [36]) or 500-fold diluted nutrient broth (NB, [37]), respectively (Table 1). The inoculum was diluted 1:1 with 2x SL or with 250-fold diluted NB resulting in final target cell count of 8.1x10^6^ CFU/surface in both evolution experiments. Surfaces were inoculated with 50 µl droplet of inoculum in case of SL and 5x2 µl droplets of inoculum in case of diluted NB (Table 1). Inoculated surfaces were transferred into a climate chamber (Climacell EVO, Memmert, USA)) and exposed for a defined time at defined climatic conditions (Table 1). Exposure was stopped by submerging each test surface to a 50 ml centrifuge tube containing 10 ml standardized toxicity-neutralizing medium (soy casein digest lecithin polysorbate broth; SCDLP [37]) and vortexing for 30 seconds at full speed. 500 μl of the harvested suspension and its 10^-1^ and 10^-2^ dilutions were bead-plated on LB agar and incubated for 40-50 h at 37°C. Lowest dilution with non-confluent colony growth was used to prepare the next inoculum as described above. Each inoculum was also stored as a glycerol at -80° C short-term to enable picking up lineages that experienced complete lethal exposure from previous exposure cycle material. The latter was only needed for copper- exposed lineages.

Mutants carrying mutations of interest were isolated directly from evolved populations or single colonies that tolerated substantially higher silver salt concentration in the minimal biocidal concentration (MBC) assays after the evolution experiment. Those isolates were subjected to whole genome sequencing as described below and included in the metal and antibiotic tolerance testing.

### Copper, silver and antibiotic tolerance

In the test formats described below bacterial sensitivity to silver, copper or antibiotic challenge was assessed in which context use of the terms resistance and tolerance needs to be clarified. Antibiotic resistance is well defined by change of minimal inhibitory concentration (MIC) exceeding epidemiologic cut-off values for general microbiology or clinical breakpoints for susceptibility testing in clinical context [38], [39]. However, use of the term in the context of other antimicrobials, including antimicrobial metals, is often confusing. For biocidal antimicrobials, killing of the microbes measured by minimal biocidal concentration (MBC) is a more relevant parameter than MIC. By this definition, metal resistance is rare, possibly due to multiple mechanisms of action as opposed to limited mode of action of most antibiotics more easily counteracted by specific defense/resistance mechanisms. To ease understanding, hereafter resistance is used for large increases in MIC values and tolerance for substantial increases primarily in MBC values without the MIC necessarily changing.

Changes in **silver and copper surface tolerance** of the evolved populations and selected isolates were tested similarly to semi-dry exposures in evolution experiments. Surfaces used and exposure times for testing populations were identical to respective evolution experiments unless noted otherwise. For testing surface tolerance of selected isolates unused metal surfaces were employed with the intent to reduce variability in viable counts but as the unused surfaces were also slightly less efficient than repeatedly used and cleaned ones, exposure times were changed to: 6 h on copper and silver in HH conditions, 35 min on copper and 4 h on silver in LD conditions.

To determine **minimal inhibitory concentrations (MIC)** of ionic silver and copper as well as selected antibiotics of the evolved populations and selected mutants compared to the ancestral strain of *E. coli* ATCC 8739 a modified standard agar dilution method [40] was used . Briefly, 2-fold dilution series of stock dilutions of respective soluble metal salts or antibiotics were suspended in 50°C Mueller-Hinton agar (MHA; Biolife MH Agar II #4017402; Biolife Italiana, Milan, Italy) in 10x10 cm square Petri dishes. To prepare the inoculum by least hindering mutant proportions in the evolved populations and avoiding competition in liquid culture, the evolved populations and the ancestral inoculum material were streaked onto LB agar from glycerol stock and incubated at 37°C for 24 h. The resulting biomass was suspended in 5 ml PBS and pipetted into clear 96-well plate in triplicates. All the suspensions were adjusted to 10^7^ CFU/ml photometrically after which 3 μl spots (3x10^4^ CFU) of each suspension was inoculated onto MHA plates supplemented with different target substance concentrations. The results were registered as presence or absence of visible growth after 24, 48, 72 and 168 h of growth to account for possible mutants with higher tolerance but lower growth rate. Only data from 24 and 48 time points are demonstrated as the results did not change with longer incubations.

To determine **minimal biocidal concentration (MBC)** of ionic silver and copper of the populations evolved on the respective metal surfaces, a drop test protocol from Suppi *et al*. [41] was used with minor modifications. Shortly, bacterial suspensions in deionized water (low-organic exposure) or organic soil load (high-organic exposure) were exposed to 2x dilution series of silver and copper salts at room temperature in static conditions 5 µl of which was drop plated to non-selective solid LB for MBC determination after 2 h and 24 h of liquid exposure. In the tests with evolved populations, bacterial suspensions from plate cultures (as described for evolution experiments) were used instead of liquid culture to least affect the mutation frequency pattern of the population. In the tests with selected mutants, an exponential liquid LB culture was used as described in Suppi *et al*. [41].

### Assessment of dihydrogen sulfide production

To assess H2S production colorimetric test based on formation of dark insoluble metal sulfides was used. For that SL medium used in high-organic humid exposure conditions (Table 1) was supplemented with 0,5 g/l (NH4)5[Fe(C6H4O7)2] as the metal ion source, 4 g/l K2HPO4 as a buffering agent, and 1,94 g/l L-cysteine as the sulfur source in final concentrations. Overnight bacterial culture in LB broth was washed twice with deionized water, diluted to OD600=0.2 and added to 2-fold concentrated sulfide detection medium (with or without AgNO3 concentration gradient) in 1:1 ratio, mixed and closed air-tight.

Formation of dark insoluble sulfides was registered qualitatively by visual inspection after static incubation in the dark at 37°C for up to 24 h.

### Assessment of indole production

To assess indole production by tryptophanase enzyme in bacteria of interest, 0.5 ml Kovac reagent (Millipore #1.09293.0100) was added to 2 ml of overnight liquid culture of bacteria in LB broth. Pinkish red colorimetric observation was considered as proof of indole production whereas yellow to brownish color was considered an indole-negative result.

### Sequencing and data analysis

The initial inoculum used to start the 1^st^ evolution cycle and harvested biomass from the 30^th^ evolution cycle of both evolution experiments was frozen at -20°C until DNA extraction with Invitrogen™ PureLink™ Microbiome DNA Purification Kit (Thermo Fisher Scientific; #A29790). All 20 populations of HH experiment, and 5 copper, 5 silver, 4 steel and 4 glass populations from LD experiment were sequenced. DNA samples were sequenced by Novogene Co. Ltd using Illumina NovaSeq 6000 resulting in 150 nt paired end reads with ∼368±167-fold genomic coverage for *E. coli* ATCC8739 out of which >98% of the sequences per sample mapped to the reference genome (GenBank: CP000946.1). Raw reads were deposited to the European Nucleotide Archive (ENA), accession PRJEB75034. Breseq analysis pipeline was used for mutation analysis in polymorphism mode [42], [43]. Mobile genetic elements in the reference sequence of the ancestor strain were detected by ISfinder [44].

## Statistical analysis

In GraphPad Prism 9 software one-way or multifactorial ANOVA with multiple comparisons at α=0.05 was used to determine statistical significance of the quantitative results. CFU values were log10 transformed prior to statistical analysis. Median values are used in the text to describe effect sizes.

## Results and discussion

### Experimental system

To illustrate the importance of exposure conditions on different evolutionary scenarios in response to antimicrobial surface exposure, *E. coli* was exposed to the antimicrobial silver and copper surfaces or stainless steel and glass control surfaces in either high-organic humid (HH) or low-organic dry (LD) exposure conditions as detailed in Table 1. These contrasting exposure conditions enabled either limited growth without complete drying of inoculum (HH) or drying of inoculum droplets and decreased viability of bacteria (LD) on control surfaces mimicking plausible spray contamination scenarios as previously described by Kaur *et al*. [2]. Cyclic transient biocidal exposure to copper and silver surfaces in LD conditions was expected to select for survival [33], [45], [46], [47] and/or more passive and less energy-expensive bacterial defense mechanisms. Cyclic bacteriostatic exposure to copper and silver surfaces in HH conditions was expected to select for growth in the presence of the antimicrobial and/or energy-expensive defense mechanisms often associated with antibiotic cross-resistance [8], [9] similarly to published examples of (sub)inhibitory ionic Ag or Cu exposures. The use of *E. coli* strain ATCC 8739 in laboratory evolution experiments was expected to offer a good model for additionally assessing the role of efflux in metal detoxification as it harbors a pathogenicity-associated [48] cryptic chromosomal copper and silver tolerance island [49], [50] with can be activated by spontaneous mutations in the *silRS* two component system derepressing expression of the SilCBA silver efflux system [50], [51].

The evolution experiments consisted of 30 cycles of exposure of *E. coli* ATCC 8739 on antimicrobial copper and silver or steel and glass control surfaces in 5 parallel lineages (Table 1, Fig 1). Between exposures the survivors were clonally grown on solid medium to increase cell count for re-inoculation of surfaces, accumulate mutations and reduce selection caused by the experimental system itself. Growth on solid medium was expected to decrease competition and selection for growth rate compared to liquid medium as mutations that increase copper surface tolerance have been indeed shown to cause slower growth [34] and extended single-cell lag phase can itself be a tolerance mechanism [53]. In fluctuating natural ecosystems trade-offs between growth and survival can be more beneficial than maximizing growth rate in optimal conditions [52]. Compared to liquid cultures where a larger proportion of survivors can be propagated and concentrated, the choice of solid medium causes stronger bottlenecking of the evolving populations which increases genetic drift. On one hand, tightly bottlenecked populations might be more prone to accumulation of high-rate weakly beneficial mutations than low-rate highly-beneficial mutations [54] but tight bottlenecking also better reflects scenarios of host-to-host transmission of pathogens [55] or microbial transfer to and from surfaces in real applications. On the other hand, selective bottlenecks have been demonstrated to lead to rapid fixation of beneficial mutations [56], [57].

**Figure 1.**
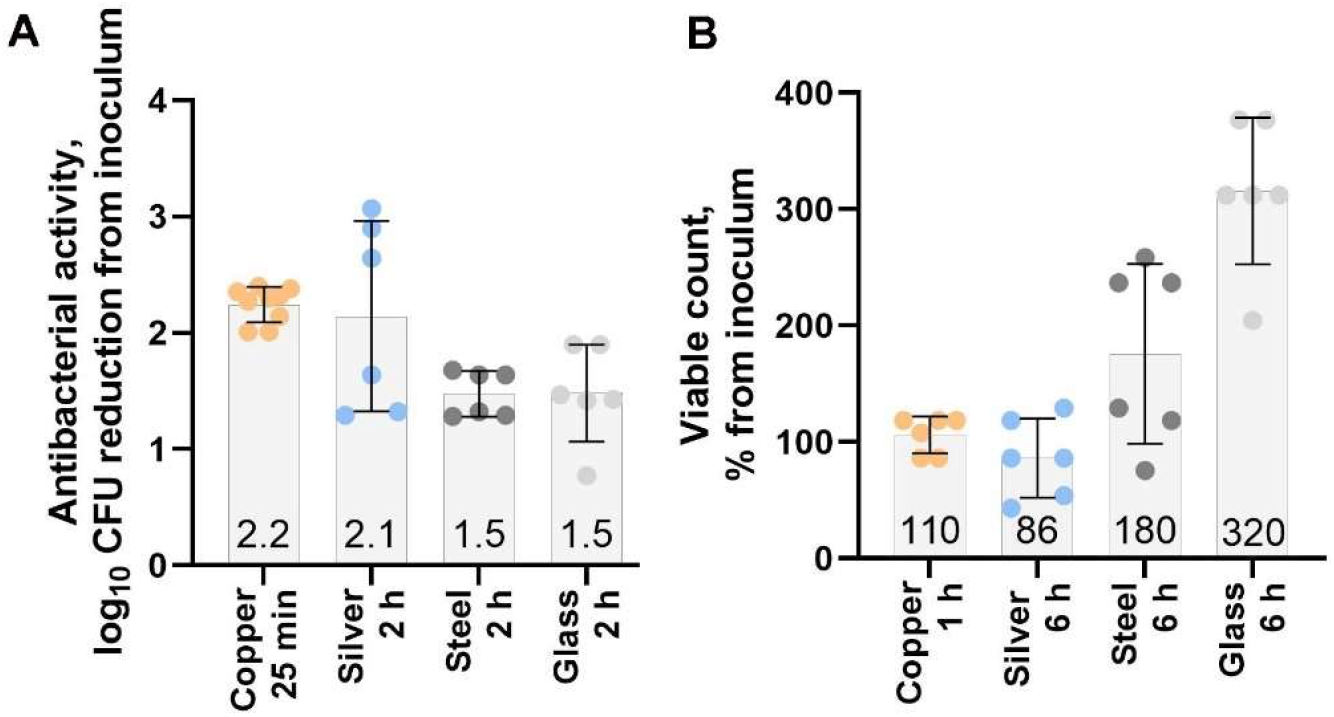
Initial bactericidal activity of the surfaces and exposure conditions in low-organic dry (LD) conditions. (A) and bacteriostatic effect of metal surfaces in high-organic humid (HH) conditions (B) against the ancestral *E. coli* ATCC 8739 strain in evolution experiment conditions. Results for antimicrobial copper and silver surfaces or stainless steel and glass control surfaces are presented. Mean values of data points from 3 independent experiments of antibacterial activity (log10CFU/surface in inoculum – log10CFU/surface post-exposure) (A) and smaller changes in viable counts relative to inoculum (%) indicating growth (B) ±SD are presented on respective bars on graph.

Due to higher toxicity of copper compared to silver and substantially different efficacy of copper and silver surfaces in LD and HH conditions as previously described by Kaur *et al*. [2], optimal exposure duration on copper and silver surfaces had to be determined for each exposure condition separately. Exposure time on control surfaces was matched to longest exposure on antimicrobial surfaces in the same exposure conditions to maximally reflect selection due to exposure conditions. The conditions selected for LD experiment aimed to kill most microbes with a target of 2-log reduction in viable counts compared to initial inoculum on antimicrobial copper and silver surfaces (Fig 1A). During the 2 h exposure, the LD exposure conditions also caused a 1.5-log reduction in viable counts on glass and steel control surfaces. Exposure conditions in the HH experiment were selected not to kill but to inhibit growth on silver and copper surfaces while limited growth with up to 180% and 320% increase in viable counts could be observed on steel and glass control surfaces, respectively (Fig 1B).

To characterize phenotypic changes and their dependance on exposure conditions, changes in silver and copper surface tolerance of the evolved populations were tested in semi-dry exposure conditions and compared to the conventional minimal inhibitory concentration (MIC) and minimal biocidal concentration (MBC) assays of the respective metal ions in liquid test format. Whole genome sequencing and mutation analysis was used to look for functionally plausible potentially beneficial mutations. To evaluate risk for the development of antibiotic cross-resistance, antibiotic MIC values of the evolved populations were measured and potential risk indicators, *e.g.* increased mutation frequency or mutations in known resistance genes and efflux systems was looked for in mutation analysis.

### Tolerance and resistance profiles of the evolved populations

Release of metal ions is considered to be one of the main biocidal mechanisms of antimicrobial metal surfaces [58], [59]. Therefore, increase in copper and silver tolerance was expected to be the most immediate result of the copper and silver surface exposure. Evolutionary changes in growth (assessed as MIC) and survival (assessed by MBC) in the presence of an antimicrobial substance might occur independently of each other and increased survival might be detected in case of unchanged MIC [45].

Furthermore, survival of metal-exposed bacteria is heavily dependent on organic content of exposure media [41], [60] and can cause up to 1000-fold differences in MBC value [41]. Therefore, we firstly assessed silver and copper tolerance of the evolved populations employing the conventional agar microdilution assay for MIC [40] and MBC assay in liquid test format [41].

Less than 2-fold changes in the ionic silver MIC values and no significant changes in ionic copper MIC values of the copper- and silver-exposed evolved populations compared to ancestor were observed (Fig 2). Also, no notable changes were observed in the MIC values of ampicillin, ciprofloxacin, gentamicin and colistin (Supplementary Fig S2) confirming lack of measurable risk of developing antibiotic cross- resistance in response copper or silver surface exposure in the conditions used. Changes in MBC values demonstrated more variability than MICs but also were small (≤2-fold increase compared to ancestor) (Fig 3).

**Figure 2.**
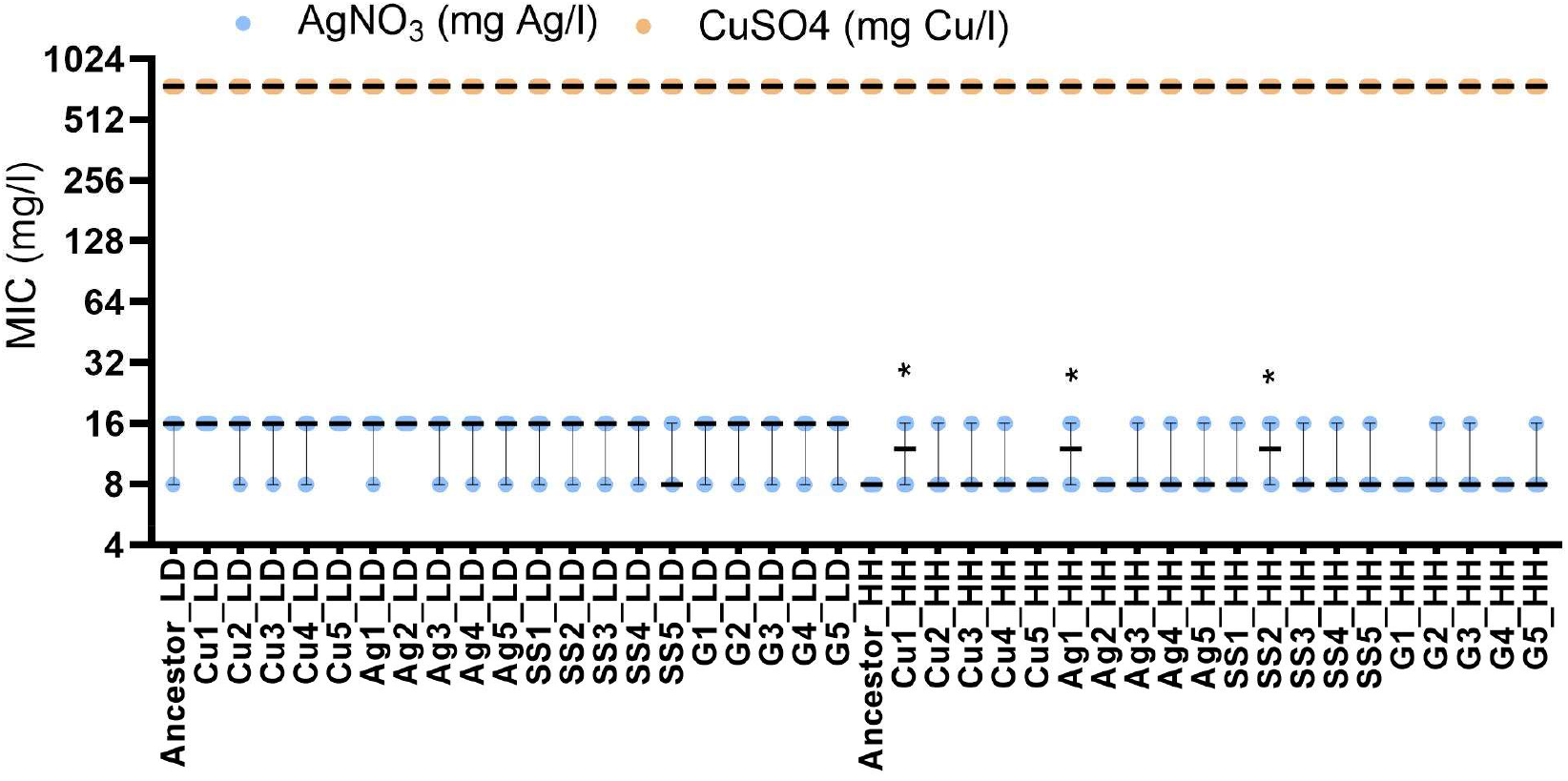
Minimal inhibitory concentration (MIC) of ionic copper and silver of the evolved populations and their respective ancestors. Copper and silver salts were used as a source of respective metal ions for MIC assay on Mueller-Hinton agar. Values from three biological and two technical replicates are presented with median and range. Statistical significance of the difference from the ancestor exposed in the same conditions is marked with * P< 0.05.

**Figure 3.**
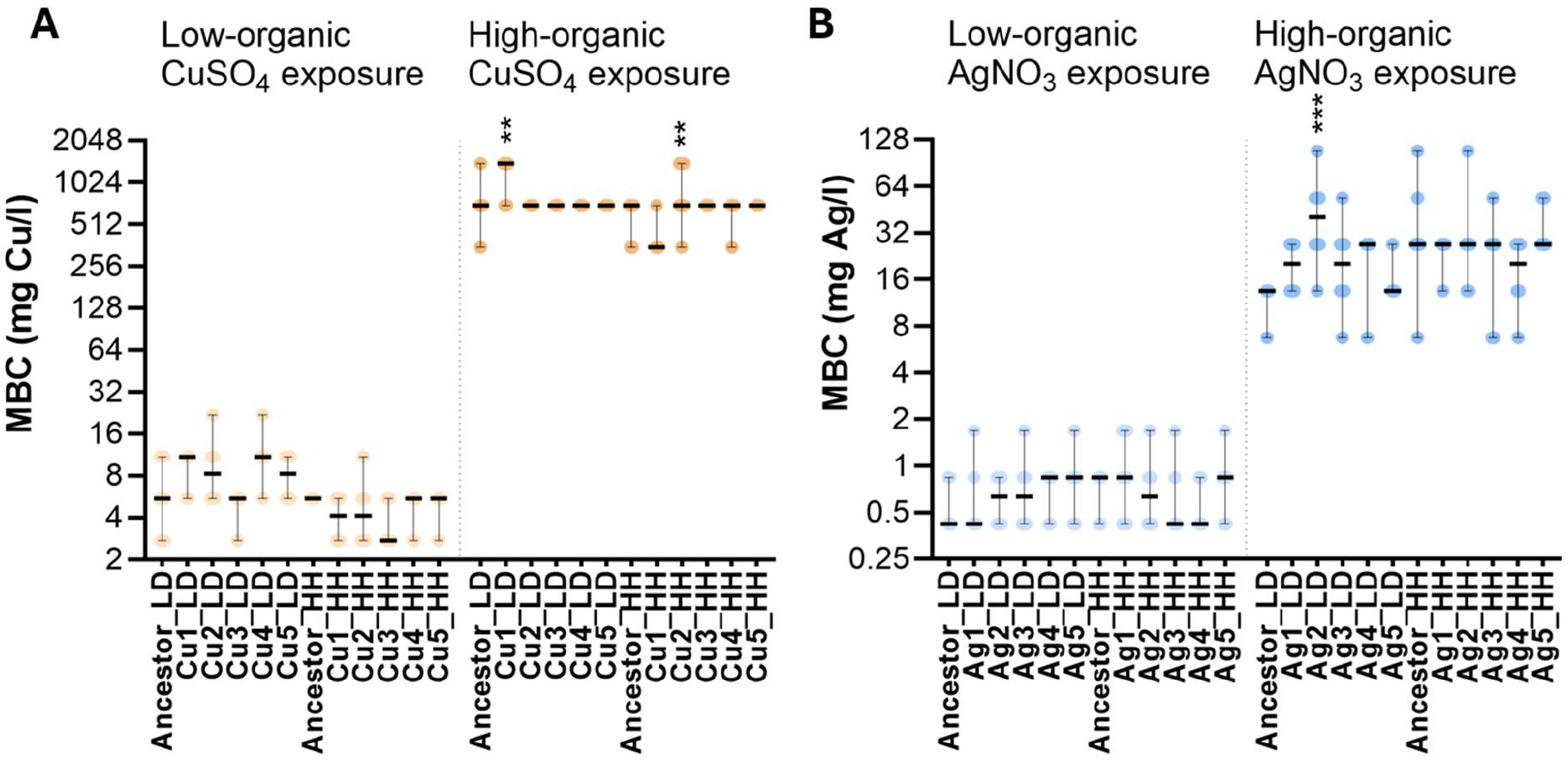
Minimal biocidal concentration (MBC) of copper and silver of the evolved populations and their respective ancestors. Populations evolved on copper surfaces were tested against copper (A) and populations evolved on silver against silver (B). Values from three independent experiments (6 data points) are presented with median and range. Statistical significance of the difference from the ancestor exposed in the same conditions is marked with ** P< 0.01; *** P<0.001.

### Silver and copper surface tolerance of the evolved populations

Evolution of tolerance can be difficult to interpret due to involvement of multiple bacterial defense mechanisms, affected by genomic and transcriptomic changes or even global stereotypic transcriptome expression patterns [61]. What is beneficial in in one specific stress condition might not be beneficial in different exposure conditions. Copper-exposed populations not being more tolerant to ionic copper is in good agreement with Bleichert *et al*. [33] who also found no increase in ionic copper tolerance of the bacteria evolved on copper surfaces despite surviving on copper surfaces for 60 min compared to <1 min of the wild-type. Xu *et al*. [34] observed 37.8% survival of copper-exposed populations on copper surfaces compared to 0.09% survival of control populations after 60 min challenge while MIC of ionic copper increased only negligibly from 3.25 mM to 3.5-3.75 mM. Also, copper tolerant bacteria isolated from dry copper alloy surfaces have demonstrated no increase in copper ion tolerance in liquid conditions [62]. In addition, we have previously demonstrated that sublethal response to copper on copper surfaces is correlated with copper content in the surface material and not the amount of copper released from the surfaces [30] suggesting a more important role of direct microbe-surface contact than the ion release. Together that might indicate that either copper and silver tolerance of the evolved populations was not changed or was changed below the detection limits of the employed methods of characterization, and/or ion release might indeed not be the main mechanism of action of silver and copper surfaces and therefore metal ion tolerance is not strongly selected during cyclic exposure to the respective metal surfaces. Furthermore, despite possible methodological challenges in detecting small changes in phenotypic traits, even a slight advantage in fitness may become biologically relevant.

Therefore, silver and copper surface tolerance of the evolved populations compared to the ancestral strain in the respective low-organic dry or high-organic humid evolution experiment was also assessed in the semi-dry evolution experiment conditions (Fig 4).

**Figure 4.**
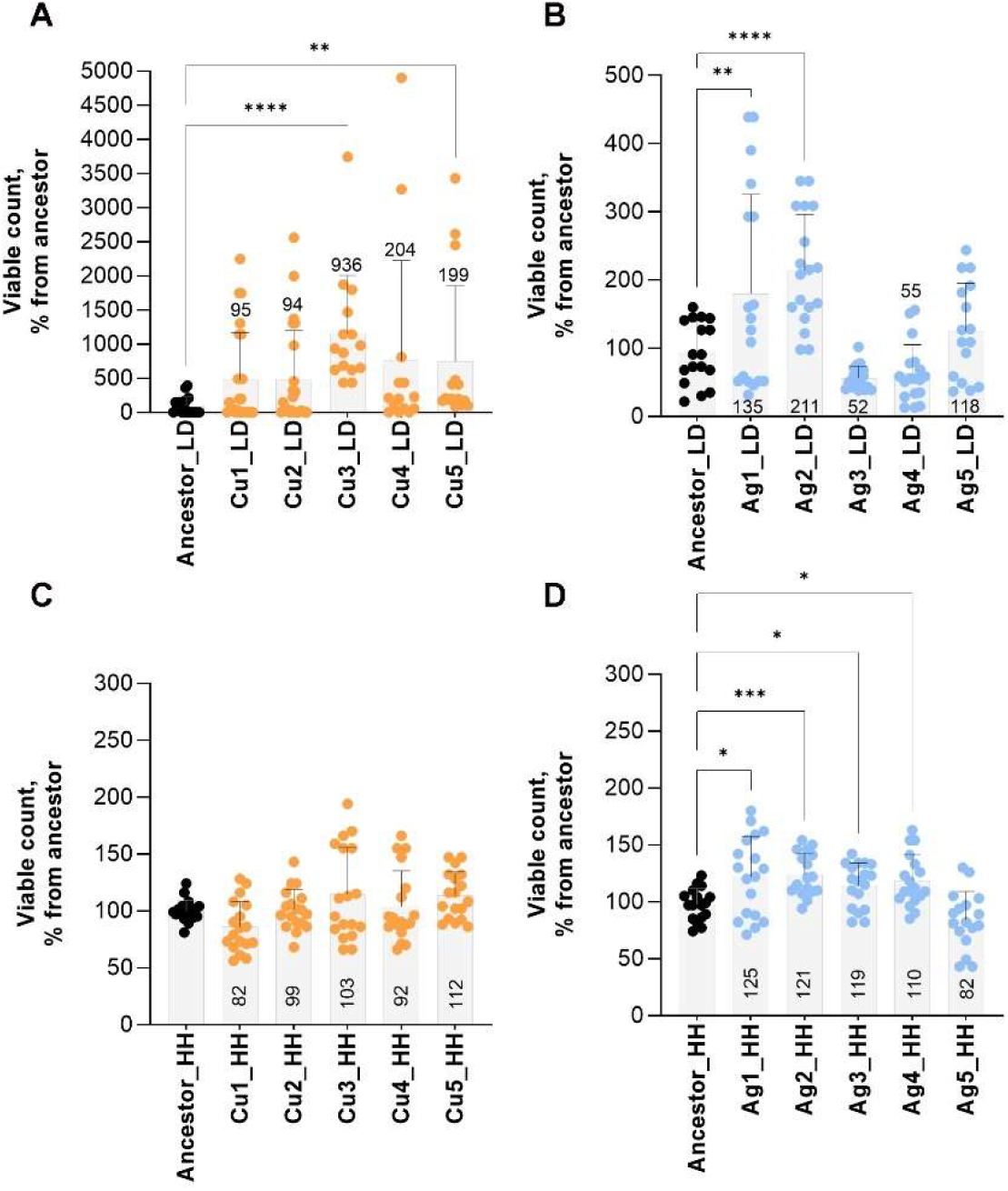
Silver and copper surface tolerance of the evolved populations. Populations evolved in low-organic dry (A,B) or high-organic humid (C,D) environments exposed to copper (A,C) or silver (B,D) surfaces were tested against the ancestral strain in the same conditions as in the corresponding evolution experiment. Values from three independent experiments are presented with mean ±SD. Median value is presented numerically. Statistical significance of the difference from the ancestor exposed in the same conditions is marked with * P< 0.05; ** P< 0.01; *** P< 0.001; ****P< 0.0001 based on ANOVA followed by *post-hoc* testing for multiple comparisons.

Largest changes in antimicrobial surface tolerance were observed for copper-exposed populations in LD conditions. Cu3_LD resulted in about 9 times and both Cu4_LD and Cu5_LD about 2 times more survivors than the ancestor (Fig 4A) and survivors were retrieved from all copper surfaces (Supplementary Fig S3). Survivor counts from Cu3_LD were at an average only 0.63 log10 decreased from the viable count in the inoculum (Supplementary Fig S3) indicating substantial increase in copper surface tolerance and survival in the LD exposure conditions compared to 2.2 log10 decrease for the ancestor (Fig 1A). Survivor counts from Cu4_LD and Cu5_LD (Supplementary Fig S3) could be considered similar to the ones of ancestor on control surfaces (Fig 1A) indicating increased copper surface tolerance in the defined exposure conditions. The Cu1_LD and Cu2_LD populations were characterized by highly variable viable counts similarly to the ancestor in the same conditions with neither presenting survival benefit over the ancestor (Fig 4A) and no survivors retrieved from several copper surfaces (better visualized on Supplementary Fig S3).

On silver surfaces in LD conditions Ag1_LD demonstrated 35% higher viable count than the ancestor mainly attributable to a few surfaces with very high survivor numbers (Fig 4B). Ag2_LD showed a more consistent about 2-fold higher viable count than the ancestor while viability of rest of the evolved Ag_LD populations on silver surfaces was not significantly different from the ancestor. The variability of viable counts on silver was generally lower than on copper and survivors were retrieved from all silver surfaces (Supplementary Fig S3).

In high-organic humid conditions none of the Cu_HH populations demonstrated significant changes in viability on copper compared to the ancestor (Fig 4C). However, Ag1_HH to Ag4_HH but not Ag5_HH demonstrated 10-25% higher viable counts on silver surfaces than the ancestor (Fig 4D). This change can seem small but could be accountable to either decrease in already small (if any) mortality or even to an extent overcoming growth inhibition on silver surface compared to control surfaces. Considering high variability and growth-supporting conditions in HH exposure, double selection for increased survival in early exposure and increased growth in later exposure could also be possible as recently demonstrated for benzalkonium chloride [63]. In either case the expected maximal possible change is low due to very limited growth potential in the exposure duration and conditions compared several logarithms of possible decrease in mortality in LD conditions.

### Mutation frequency and types of mutations

In addition to changes in antimicrobial tolerance, our aim was also to assess adaptation potential and possibly increased risk of AMR development. One of the indicators of quicker adaptation and increased risk of AMR development and virulence is the emergence of hypermutators resulting in increased mutation frequency. In whole genome sequence data of populations evolved on copper, silver or steel surfaces no significant differences in the number of accumulated single nucleotide variants (SNVs) compared to populations evolved on biologically inert glass control surface were found (Fig 5A). This result opposes the finding by Xu *et al*. [34] who found increased mutation accumulation in *P. fluorescens* evolved on copper surfaces. Although Xu *et al*. explained the high mutation frequency with *mutL* mutations in copper-exposed lines, they detected the *mutL* 1174_(G)5 -> (G)4 change as well as several other frequency-matched SNVs in other genes in all eight parallel copper-exposed populations that could indicate same ancestral origin of the mutations and not selective enrichment of parallel mutations in *mutL* on copper surfaces. There were no mismatch-repair system mutations present in copper or silver exposed lineages in our results. The conflicting results highlight the need to use the same inoculum for test samples and controls and to sequence the material from the first inoculum of the evolution experiment.

**Figure 5.**
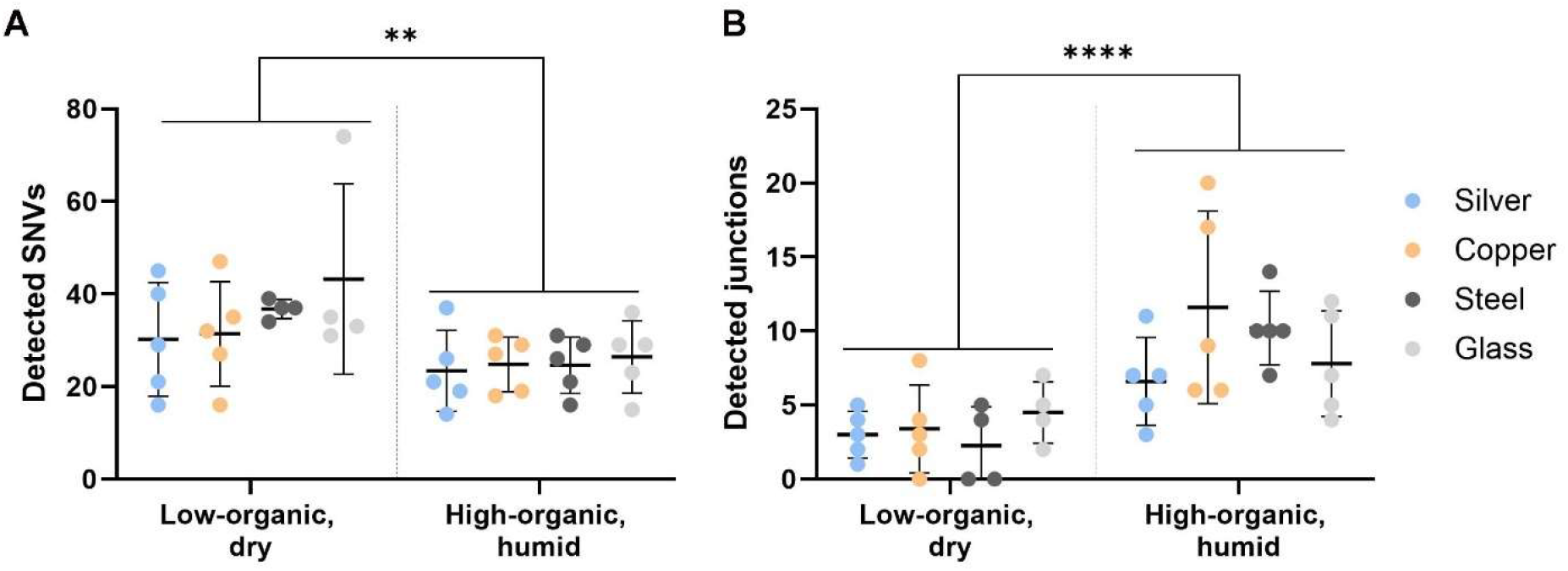
Total number of single nucleotide variations (SNVs) (A) and larger genomic rearrangements (new junctions) (B) detected in the whole genome sequencing (WGS) data of the populations evolved on antimicrobial silver and copper or steel and glass control surfaces. Total number of single nucleotide variations (SNVs) and total number of larger genomic rearrangements (new junctions) detected in the WGS data of the evolved populations from high-organic humid (HH) and low-organic dry (LD) exposure conditions at ≥0.05 frequency after removing mutations present in the ancestor. There were no significant differences in the number of SNVs (A) or junctions (B) accumulated during the evolution on metal surfaces compared to the control evolved on glass in the same conditions. Significantly higher number of SNVs were detected in LD exposure conditions compared to HH in general with the exposure environment explaining 23% of total variation in SNV counts and 49% of total variation in transitions to transversions ratio based on two-way ANOVA with post-hoc testing for multiple comparisons. Several larger genomic arrangements were detected (B), usually involving IS element mobility and rDNA reorganization, but as not all such events could be fully annotated using short-read data, total number of new sequence junctions detected is presented. Significantly higher number of junctions were detected in HH exposure conditions compared to LD in general with the exposure environment explaining 41% of total variation in junction counts based on two-way ANOVA with post-hoc testing for multiple comparisons. The ancestral inoculum of the HH experiment also contained more rDNA reorganization events compared to the reference genome than the one of the LD experiment and could lead to a random bias. However, the same association held after removing all rDNA reorganizations from the comparison, with exposure environment still explaining 20% of total variation (Supplemetary Fig S4). ** P< 0.01; **** P< 0.0001

The types of substitutions among SNVs can be indicative of selection pressure and the mechanism of action of the antimicrobial used. Among the detected SNVs, there were no significant differences in the ratio of transitions to transversions in the populations evolved on metal surfaces compared to glass (Fig 6A). However, generally higher than the expected (*e.g.* 44% transversions in wild-type *E. coli* MG1655 [64]) transversion accumulation that is not contributable to oxidation-prone G>T substitution (Fig 6B) but rather A:T>T:A (Supplementary Fig S5) was unexpected. Similar tendency on all surface types suggests a strain-specific effect. Unexpectedly high presence of transversions and even higher transversion proportion in LD conditions than HH conditions could indicate nutrient limitation [65] and stronger purifying selection [66]. High level of T:A>A:T substitutions is scarcely described in previous studies but has been indicated in highly transcribed regions [67] or in case of specific mutants *e.g. recA*730 (E38K) [68]. Higher proportion of G->T substitutions was found on copper compared to glass in high-organic humid conditions (Fig 6B) possibly indicating oxidative damage of guanine [69], [70].

**Figure 6.**
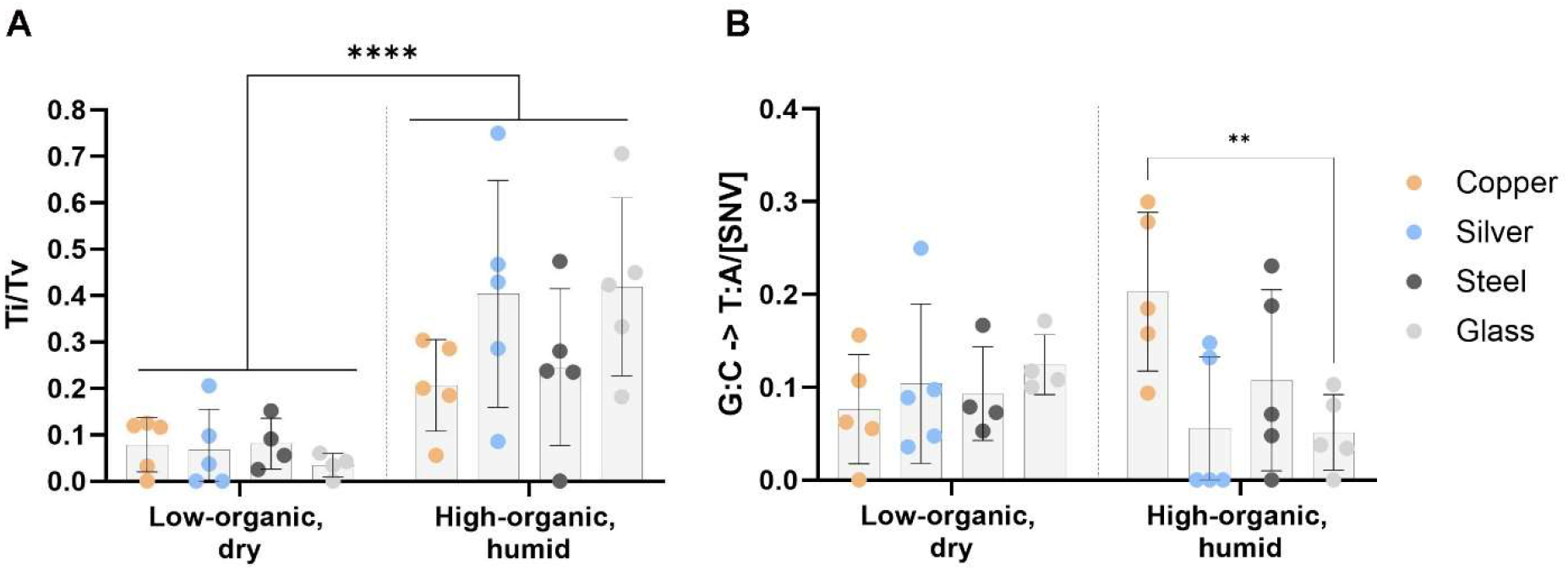
Types of single nucleotide variations (SNVs) detected in the populations evolved on antimicrobial silver and copper or steel and glass control surfaces. Proportion of transitions (Ti) to transversions (Tv) among the detected SNVs (A) and guanine to thymine substitution proportion among the SNVs (B) detected in the WGS data of the evolved populations from high-organic humid (HH) and low-organic dry (LD) exposure conditions at ≥0.05 frequency after removing mutations present in the ancestor. There were no significant differences in transitions to transversions ratio (A) accumulated during the evolution on metal surfaces compared to the control evolved on glass in the same conditions. The environment of the evolution experiments (HH vs LD) did not contribute to variation of G->T substitutions (B) but compared to glass more G->T transversions were accumulated on copper in HH conditions. ** P< 0.01; *** P< 0.001

Reactive oxygen species (ROS) damage on copper surfaces is an expected mechanism due to copper [71] and iron [72] producing Fenton-like reactions unlike silver and glass.

In addition to SNVs larger genomic rearrangements (Fig 5B), especially IS1-mediated deletion enrichment was abundantly observed after exposure to control surfaces in high-organic humid conditions where bacteria were proliferating also during surface exposure and were much less frequent, but not missing, on antimicrobial surfaces in low-organic dry conditions (Fig 5B; Table 2; Supplementary Table S1). The initial inoculum of the high-organic humid experiment also contained more rRNA reorganizations than the one for LD experiment (Fig 5B; Supplementary Fig S4). The cause of this variation as well as its possible effect on evolutionary scenarios remains unknown.

**Table 2.**
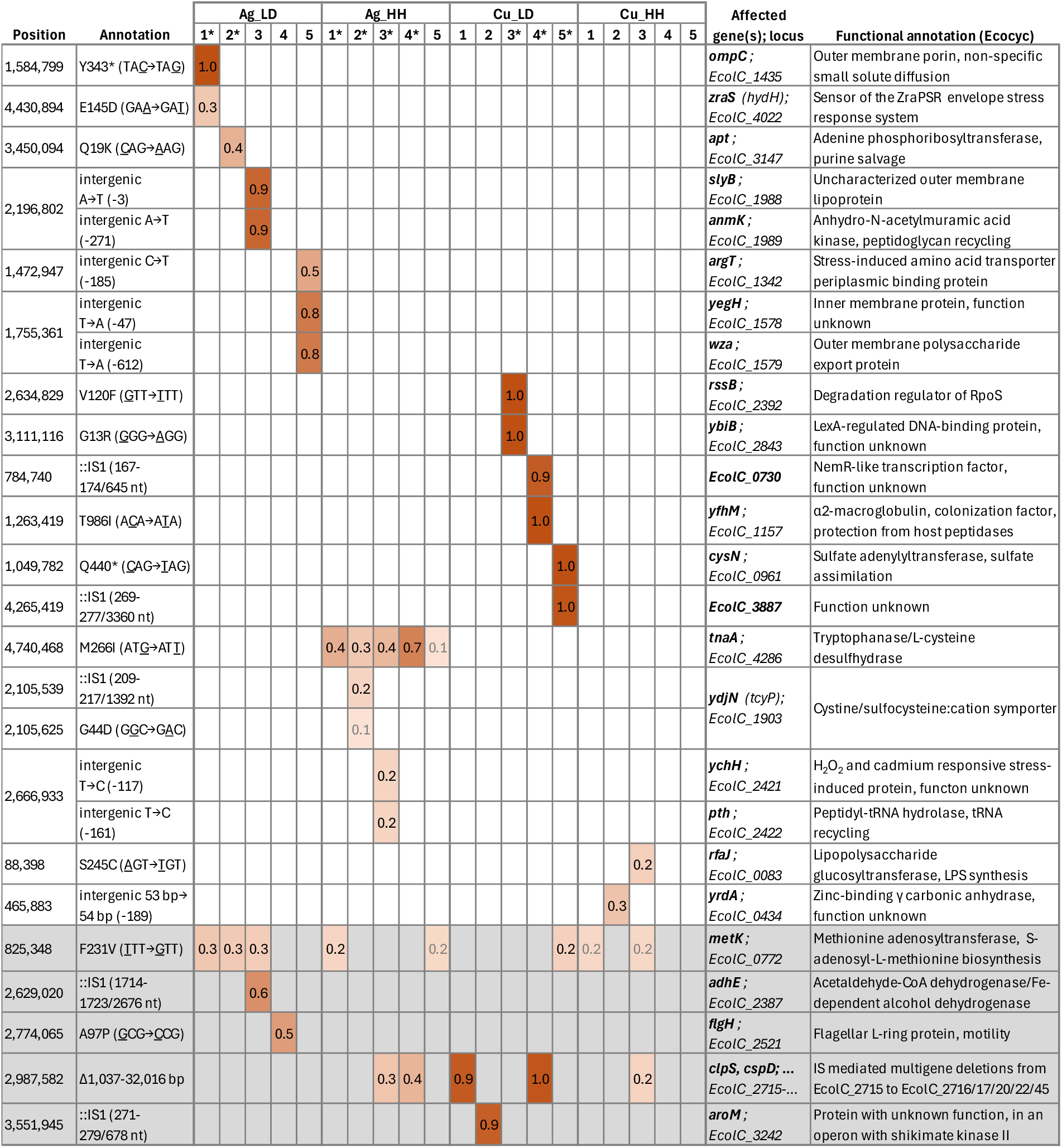
Mutations accumulated in populations evolved on copper and silver surfaces in low-organic dry (LD), and high-organic humid (HH) exposure conditions at ≥0.20 frequency. Populations with significantly increased metal surface tolerance are marked with an asterisk int the table header. Grey numbers indicate mutation frequency <0.20 in cases where mutations affecting the same gene also occur at ≥0.20 frequency in other populations. Grey background marks the genes with mutations accumulated also on control surfaces. Synonymous SNVs, rRNA reorganizations and intergenic changes downstream of neighboring reading frames are excluded from the table. Full mutational profiles of all ancestors, evolved populations and isolates can be found in Supplementary Results file.

IS element presence in non-pathogenic bacterial strains is well known [73]. In the genome of *E. coli* ATCC 8739 forty-eight IS elements from eight IS families were detected with IS1 elements being the most abundant (Supplementary Fig S6). Deletions directly adjacent to existing IS1 elements could possibly be explained by abortive transposition of the IS resulting in flanking deletions of variable length [74].

Similar phenomenon has been recorded for example in previous *E. coli* evolution experiments that have resulted in loss of motility due to deletions in flagellar genes next to an IS1 element [75] and in case of deletions next to IS1X2 element causing adaptive nitrofurantoin resistance in pathogenic *E. coli* [76].

IS1X2 has also been found to be significantly associated with antibiotic resistance genes (within 10 ORF distance) in metagenomic datasets [77].

Our results showed that IS1X2 located at *EcolC_2714* in the reference genome seems to be especially prone to cause deletions of variable length starting with *EcolC_2715.* These deletions were detected in different exposure conditions all spanning at least the reading frames of *EcolC_2715*(*clpS*) and *EcolC_2716*(*cspD*) with the longest deletion extending to *EcolC_2749*. Deletions with different lengths can be present in the same evolved population at high frequencies. Potential benefit of the deletions is not directly clear but could be speculated to be associated with *cspD* replication inhibitor inactivation rather than *clpS* because the *EcolC_2714* IS1 itself is an insertion into *EcolC_2713*(*clpA*) expected to inactivate the ClpSAP-dependent protein degradation pathway.

### Ancestral mutations

Due to separate single-colony origin of the ancestral culture of the HH and LD evolution experiments the ancestral genetic context slightly differed for the two evolution experiments as detailed in Supplementary Table S2 and Supplementary Results file. The ancestor of the HH experiment carried >80% fixed mutations in *fliI* flagellar ATPase and *oppA* genes rendering the resulting populations immotile and possibly interfering with periplasmic oligopeptide sequestering and transport. The *fliI* and *oppA* mutations in Ancestor_HH had no effect on copper (Fig 2 and 3) nor antibiotic tolerance (Supplementary Fig S2) but could have a minor effect on silver tolerance as demonstrated by 2-fold higher MBC values on Fig 3 and 2-fold lower MIC values on Fig 2 compared to Ancestor_LD. Neither of the mutations in HH ancestor significantly affected survival or growth on silver and copper surfaces (Supplementary Fig S7). However, different genetic context of HH and LD ancestors could have affected the evolutionary scenarios observed during the evolution experiment.

### Mutations in copper and silver surface-exposed populations

Data from mutation analysis was further used to identify genetic changes in known tolerance mechanisms as well identify potential new mechanisms underlying the phenotypic changes in silver and copper tolerance (Fig 4). No mutations in the genes and their regulatory regions known to be responsible for copper and silver homeostasis or granting antibiotic resistance were detected. Specific mutation accumulated on silver and copper surfaces at over 0.2 frequency in evolved populations are presented in Table 2. While trying to connect functional annotations of the mutated genes in the evolved populations (Supplementary Fig S8) to their role in (metal) stress tolerance, one must keep in mind that firstly, not all mutations in a population have been selected for and secondly, phenotypic change resulting from a beneficial mutation is not necessarily large. Functional enrichment analysis or even simpler functional clustering of mutation data with a few high-frequency mutations and several lower-frequency mutations per population suffers from high amount of non-causal functional noise and cannot be used for drawing conclusions as seen also on Supplementary Fig S8. Random mutations in the same genome with a beneficial mutation enrich in frequency just by genetic linkage (“hitchhiking”).

Therefore, although the mutations enriched with the highest frequency in any given population are the most probable causal genetic change that increases fitness in the selective context, but the functional causality of the mutations needs to be further tested by phenotypic assessment of single gene mutants as opposed to heterogenic populations or isolates from these populations inevitably carrying more than one mutation (*e.g.* isolates in Supplementary Table S2).

### High-frequency mutations in copper surface-exposed populations with increased surface tolerance in low-organic dry conditions

Largest changes in antimicrobial metal surface tolerance (Fig 4) as a result of cyclic exposure were detected on in Cu3_LD, Cu4_LD and Cu5_LD populations evolved on copper surfaces in low-organic dry conditions. However, none of the potentially causal mutations (Table 2) significantly affected copper toxicity in conventional MIC (Fig 2) and MBC (Fig 3) assays.

The Cu3_LD population with the largest increase in viability on copper surfaces carried fixed missense mutations in genes ***rssB*** (V120F in the response regulator domain) and ***ybiB*** (G13R in glycosyl transferase domain). Function of both genes is associated with stress tolerance. RssB is a σ^S^ stability modulator in *E. coli* [78] absence of which increases σ^S^ stability and bacterial osmotic, oxidative and heat stress tolerance [79]. YbiB is a poorly characterized LexA-regulated DNA-binding protein that might be involved in SOS response and persistence [80].

Cu4_LD had a fixed IS-mediated multigene deletion, the kind of which were also present on control surfaces and is not expected to increase copper tolerance. It also carried a fixed SNV in ***yfhM*** (T986I) and high-frequency (0.9) IS1 insertion into a reading frame (EcolC_0730) encoding a nemR-like transcriptional regulator. YfhM is a colonization factor, an α2-macroglobulin that traps host proteases and works in close association with peptidoglycan glycosyltransferase acting as a as a defense and repair mechanism [81], [82].. Copper toxicity, has been previously associated with envelope stress, impaired lipoprotein maturations and peptidoglycan cross-linking [6].

Cu5_LD also carried a fixed IS1 insertion in the reading frame ***EcolC_3887*** with unknown function and premature stop codon in ***cysN*** (Q440*) both possibly impairing the function of the encoded proteins. CysN is one of the first enzymes in cysteine biosynthesis pathway by activating sulfate in sulfur assimilation process. Interestingly, up-regulation of cysteine biosynthesis genes, including *cysN*, has been indicated in response to metal ions above physiological concentrations, *e.g.* zinc [83] and copper [84]. Furthermore, SNVs in *cysN* of *P. fluorescens* were also fixed in 2 out of the 8 copper-exposed populations in the evolution experiment by Xu *et al*. [34].

### High-frequency mutations in silver surface-exposed populations with increased surface tolerance in low-organic dry conditions

On silver surfaces significantly increased survival was detected in Ag1_LD and Ag2_LD populations (Fig 4) while only Ag2_LD demonstrated a slightly increased copper tolerance in MBC assay (Fig 3) but only in high-organic conditions and neither population had a different MIC value (Fig 2) from the ancestor.

The Ag1_LD carried a fixed mutation in ***ompC*** (Y343*) expected to impair the encoded protein and a lower-frequency (0.3) substitution in ***zraS*** (E145D). The outer membrane proteins OmpC and OmpF, are the two major porins of *E. coli*. Although cross-membrane transport and tolerance mechanisms of silver and copper ions in bacteria partly overlap, it is known for decades that conditions selecting for silver tolerance might not by default also grant increased copper tolerance [7]. The role of outer membrane porins in metal tolerance is somewhat unclear. For example, it has been demonstrated that the lack of *ompC* and/or *ompF* alone does not seem to grant significantly increased silver and/or copper salt MICs in *E. coli* laboratory strains [7] while deletion of one or both of the genes from *E. coli* MC 4100 increased tolerance to both silver ions and nanoparticles [85]. It can also be that while defective porin function alone is not enough then mutations in both passive influx by Omp porins and active efflux by Cus can increase silver tolerance of *E. coli* [48], [51], [86]. Complexity and differences in transcriptional regulation and presence or absence of functional homologues in different *E. coli* strains might influence individual porin role in metal tolerance. Although mutations in *ompC* have been associated with antibiotic resistance [87], [88], MIC values of the populations with fixed *ompC* mutation did not demonstrate changes in ampicillin, ciprofloxacin, gentamicin or colistin MIC values (Supplementary Fig S2).

There seems to be confusion about the *ompC* genes in the published *E. coli* genomes. The *ompC* gene seems to have undergone substantial mosaic evolution [89] and *ompC* of *E. coli* ATCC 8739 has been previously described to be disrupted by an IS1 insertion [90]. Surprisingly, neither o*mpC* (*EcolC_1435*) of the *E. coli* ATCC 8739 reference genome (CP000946.1)[90] nor respective genes in other accessible ATCC 8739 genomes (CP022959.1; CP033020.1) had a truncated *ompC*. The *E. coli* ATCC 8739 sequence CP043852.1 seemed to have a 100% similar *ompC* to K-12, lacked the *pco* and *sil* gene clusters and the genome might be wrongly designated to ATCC 8739 strain. Resequencing of *the E. coli* ATCC 8739 ancestor strain revealed no deletions or IS insertions in *ompC*. Therefore, we expected *E. coli* ATCC 8739 to have a full length functional *ompC*. Furthermore, as described above, we detected a fixed mutation of a premature stop codon (Y343*) in *ompC* in the silver-exposed Ag1_LD line indicating selective pressure towards *ompC* inactivation.

ZraS E145D in histidine kinase of the ZraSR two-component envelope stress signaling pathway that was enriched in Ag1_LD population could be a hitchhiker to a fixed OmpC Y343* in the same population but could also plausibly contribute to metal tolerance. Similarly to *ompC*, *zraS* is also indicated in the Gram-negative envelope stress response via CpxAR-regulation [91] and their individual or possibly synergistic roles in silver tolerance should be studied further. The observed amino acid substitution occurs in the periplasmic sensor-like domain (InterPro: P14377 [92]), not the kinase domain. Mutations in sensory domain of ZraS might change sensor affinity to other metals besides the already known Zn and Cu [93].

More interesting might be the mutation with the highest frequency in the Ag2_LD population with 2- fold more survivors than ancestor on silver surfaces in low-organic dry conditions (Fig 4) as well increased MBC in high-organic exposure but not MBC in low-organic exposure (Fig 3) or MIC value (Fig 2). Ag2_LD carried no fixed mutations, but a 0.4 frequency substitution in ***apt*** (Q19K, N-terminal from the phosphoribosyltransferase domain). It is not immediately clear how mutations in adenine phosphoribosyltransferase *(**apt**)* could increase silver surface tolerance. If the mutation impairs purine salvage and adenine is not converted to AMP by Apt or the reaction is less efficient, one could suspect that as adenine has high affinity towards silver ions [94], [95], some of the adenine that is not recycled or is processed by Apt less efficiently could in theory participate in silver detoxification intracellularly and possibly also extracellularly. Disruption of *apt* has also been shown to cause reprogramming of nucleotide metabolism, adenine accumulation and increased copper tolerance while not affecting intracellular copper accumulation in *S. aureus* [96]. Furthermore, overexpression of not only *apt* but also other enzymes that use phosphoribosyl diphosphate (PRPP), sensitized *cop*^-^ strain of *S. aureus* to Cu^2+^ indicating a more general involvement of Cu^1+^ induced blocking of the pentose phosphate pathway and sparing of PRPP protecting against Cu^2+^ toxicity. As toxicity, transport and detoxification of Ag^1+^ largely overlap with Cu^1+^ it is plausible that possible impairment of *apt* offers protection against both.

The mutations carried by Ag3_LD – Ag5_LD, although some of them in genes involved in stress responses (*argT*), possibly affecting cell surface properties (*wza*, *anmK*, *slyB*) or transport function (*yegH*), do not measurably change metal surface tolerance which does not rule out possible benefit, but multiple mutations could be needed to achieve a substantially more tolerant phenotype.

### High-frequency mutations in copper or silver surface-exposed populations with increased surface tolerance in high-organic humid conditions

No changes in copper tolerance of the populations exposed to copper compared to ancestor were observed in high-organic humid conditions possibly due to weak selection. The relatively low-frequency mutation in ***yrdA*** (0.3 in Cu2_HH) and ***rfaJ*** (0.2 in C3_HH) might potentially have been enriched due to selection but in that case their phenotypic effect and/or frequency is too small to cause significant changes in viable counts. *RfaJ* participates in lipopolysaccharide (LPS) synthesis [97]. Interestingly, both *rfaJ* mutated in Cu3_HH in our study and mutated *rfaP* of copper-exposed *P. fluorescens* in Xu *et al*. study seem to be involved in the biosynthesis of the core oligosaccharide region of LPS. *YrdA* with an upstream (−189) mutation in Cu2_HH on the other hand seems to be encoding a cytosolic Zn^2+^-binding hexapeptide repeat acetyltransferase, gamma carbonic anhydrase like protein with unknown function that is induced by polyamine and azide [98], [99], [100] and is possibly associated with assimilatory sulfate reduction [101].

More notable mutations in genes associated with sulfur metabolism were detected in silver-exposed populations in high-organic humid conditions. M266I substitution in tryptophanase (***tnaA***) was enriched in all silver-exposed populations (frequency 0.13-0.70) and not on any other surface or in low-organic dry conditions strongly indicating enrichment due to selection. Synonymous SNVs with matching frequency to *tnaA* M266I in *EcolC_3078* and *EcolC_3609* reading frames in all Ag_HH populations (Supplementary Results for more detail) indicate selective enrichment of a low-frequency ancestral mutation. The populations with higher *tnaA* mutation frequency (Table 2) also demonstrated slightly increased silver surface tolerance (Fig 2) but no changes in MIC (Fig 2) or MBC (Fig 3) values.

Contribution of adaptive *tnaA* mutations to increased stress tolerance has been described before *e.g.* for heat shock but the underlying mechanisms can be challenging to interpret [102].. In the case of M266I, the conserved Ag-vulnerable methionine was substituted with structurally similar but less vulnerable isoleucine in close proximity of pyridoxal-phosphate modified K270 in the active center of the tetrameric enzyme [103]. Tryptophanase is best known for degrading exogenic L-tryptophan resulting in production of indole but also participates in detoxifying excess amounts of exogenous L-cysteine, is highly inducible by L-cysteine and well known to catalyze reactions leading to hydrogen sulfide production from cysteine [104], [105], [106]. Both the wild-type *E. coli* ATCC 8739 and its M266I mutant isolate produce indole and show increased hydrogen sulfide production in the presence of silver (Supplementary Fig S9) indicating that the mutation does not inactivate tryptophanase. Therefore, we hypothesize that the mutation could protect the enzyme function under silver stress. While indole can play a role in stress signaling inside and between bacterial cells hydrogen sulfide can freely cross bacterial membranes by passive diffusion and potentially directly detoxify intracellular and extracellular silver ions by forming insoluble silver sulfide. Out of the several enzymes capable of H2S production in *E. coli* [105], [108], [109], TnaA can be the main producer in rich medium [110]. Signaling and stress-related role of H2S in bacteria is poorly described but increased H2S production has been previously associated with increased silver tolerance of *E. coli* [111], alleviating heavy metal toxicity to fungi [111] and offering protection against oxidative stress and antibiotics [113].

In Ag2_HH population with relatively lower TnaA M266I frequency (0.29) we also observed IS1X2 insertion into ***ydjN*** (0.21) reading frame and a G44D substitution in the same gene (0.12) possibly indicating competition of beneficial mutations in the population. *YdjN* encodes one of the two cystine importers in *E. coli* [113], [114] which can import cystine in oxic conditions. YdjN is required by *E. coli* for growth on S-sulfocysteine as a sulfur source [116], has been demonstrated to contribute to oxidative stress tolerance [117], and its expression can be upregulated by silver nanoparticle exposure [118].

However, defective YdjN -mediated transport has been observed in case of *E. coli* resistant to toxic selenocystine [114]. In our case possible *ydjN* inactivation could avoid uptake of Ag-cysteine complexes in high-organic conditions in the presence of peptides and amino acids free to react with silver in the extracellular space.

To test the potentially causal roles of *tnaA* and *ydjN* mutations in tolerance profiles of *E. coli*, isolates carrying the mutations were isolated from Ag4_HH and Ag2_HH populations, respectively. Other mutations present in the isolates are described in Supplementary Table S2 and in the Supplementary Results file.

Possible enhancing effect of *tnaA* M266I in silver and copper or antibiotic tolerance was not confirmed (Supplementary Figures S10, S11 and S12). This could either mean that the potential benefit is too small to reliably detect with the methods of choice or that the accompanying mutations in the isolate could negatively compensate the mutations beneficial effect. The latter could be suspected as mutations similar to the IS-mediated deletion in *tnaA* M266I are much more frequent in the populations evolved on control surfaces (Supplementary Table S1) than on copper and silver surfaces (Table 2) which could be indicative of a mutational hotspot with negative selection on copper and silver surfaces, not positive selection on control surfaces. A single gene mutant should be constructed or a set of isolates carrying tnaA M266I to further study the role of *tnaA* in silver tolerance. However, protective role of *ydjN*::IS1 was confirmed by testing the isolate carrying the mutation. *ydjN*::IS1 isolate was significantly more tolerant to copper and silver surfaces in high-organic humid but not low-organic dry conditions compared to the wild-type strain (Supplementary Fig S10) and demonstrated higher MBC values of copper in high-organic exposure, but not low-organic exposure (Supplementary Fig S11). Copper, silver and antibiotic MIC values (Supplementary Figures S11 and S12) as well as silver MBC (Supplementary Fig S11) values of the mutant remained unchanged compared to the wild-type strain.

### Possible effect of the exposure media

The environment of the high-organic humid experiment could explain some of the context of mutations accumulated that were associated with sulfur and amino acid metabolism. The high-organic soil load (2.5 g/L BSA, 3.5 g/L yeast extract, 0.8 g/L mucin) is rich in protein with strong overrepresentation of sulfur-containing amino acid cysteine, the substrate of TnaA for H2S production and transport target for YdjN. Co-incidentally also mutations in other sulfur metabolism genes (*dcyD* in high-organic and *metK* in both exposure conditions, Supplementary Table S2) were accumulated on control surfaces. There are 35 cysteines out of 607 amino acids (5.8%) in BSA (Uniprot: P02769) and 118 cysteines out of 1390 amino acids (8.5%) in mucin (Uniprot: A0A5G2QSD1) compared to around 1% in *E. coli* in general [119]. Serum albumins are also well known to bind metal ions, metal complexes and metallic nanoparticles, including those of copper and silver [120], [121], [122] which causes structural changes in the protein and Cu(II) could cause BSA to aggregate [121]. Mucin is also known to act as a copper chaperone that binds copper in both its Cu(I) and Cu(II) forms [123]. The high cysteine content of the organic soiling medium could partly explain enrichment of sulfur metabolism mutations and should be considered while comparing the results to other exposure media. Therefore, we propose that the benefit of the sulfur metabolism associated mutation in silver and copper exposed populations was probably dependent on both the exposure medium and specific environmental.

### Ionic silver exposure in liquid, but not exposure to solid silver surface, rapidly selects for silver resistance

*E. coli* ATCC 8739 contains chromosomal ***pco*** (*EcolC_3410-3417*) copper resistance and acquisition [124], [125], [126] as well as ***sil*** (*EcolC_3420-3428*) silver resistance [51], [86], [127] gene clusters forming a heavy metal homeostasis/resistance island [49]. The whole *sil* and *pco* region shows 97% sequence similarity to the historical metal resistance plasmid pMG101 (GenBank: NZ_JAFFIC010000003.1) and could be of possible plasmid origin via an independent transposition event [128], [129]. The *sil* locus can be activated by spontaneous mutations in the *silRS* two component system derepressing expression of the SilCBA silver efflux system [50], [51].

Surprisingly, no changes in the *sil* locus of the silver-exposed populations were detected after the evolution experiment although the importance of such mutations in increased silver tolerance in addition to the mere presence of the *sil* genes have been described [50], [130] and increased tolerance resulting from *silS* mutation has been observed after only a single high-dose silver exposure [51]. Also, no mutations in known copper uptake or resistance genes were detected in populations evolved on neither copper nor silver surfaces.

Despite not observing mutations in *silRS* and no substantial changes in MBC and MIC values of the populations evolved on copper and silver surfaces, we occasionally observed single colonies growing after several folds higher dose of silver in the high-organic MBC exposure than the median values presented on Fig 3. From those post-evolution MBC assays single colonies with seemingly substantially higher Ag tolerance were isolated and retested to detect isolates with reproducibly increased metal ion tolerance. Such secondary isolates demonstrating persistent increase in copper and silver ion tolerance were sequenced and found to carry independent *silS* or *silS* and *cusA* mutations (mutational profiles described in Supplementary Table S2 and Supplementary Results file). None of these mutations were detected in their population of origin (Ag5_LD and Ag4_HH) and must have independently emerged after the evolution experiment indicating rapid selection of *silS* mutants in high-dose silver exposure in liquid exposure, but not on silver surfaces. This could be considered as indirect evidence of the following. Firstly, that metal ion efflux does not play a central role in antimicrobial metal surface tolerance and resistance in semi-dry conditions. Secondly, that ion release might not be the main mechanism of action of antimicrobial metal surfaces in semi-dry conditions.

The secondary isolates carrying *silS* mutations did not demonstrate substantial advantage during exposure to copper and silver surfaces (Supplementary Figure S10). However*, silS* mutants were at least 8 times more resistant to silver (grew at highest tested MIC concentration; Supplementary Figure S11) and demonstrated significantly higher MBC values for both silver and copper salts compared to wild-type strain. The *silS* mutants were also less sensitive to ampicillin and ciprofloxacin than the wild type (Supplementary Fig S12). *Sil* mutants have also previously been demonstrated to be less susceptible to ampicillin and ciprofloxacin [50]. This highlights the critical importance of exposure conditions not only in antimicrobial surface efficacy testing but also in tolerance mechanisms and risk assessment.

## Conclusion

In this study we were interested in phenotypic and genotypic changes in bacterial populations evolved on antimicrobial copper and silver surfaces, and the role of such changes in developing metal tolerance and antibiotic cross-resistance. As a result of evolution experiments with cyclic exposure of *E. coli* ATCC 8739 to copper and silver surfaces in semi-dry conditions metal surface tolerance was increased in some but not all evolved populations. Increased copper or silver surface tolerance was mostly not associated with increased MIC and MBC values of ionic copper and silver indicating central role of contact-killing and not ion release as mechanism of action of the respective surfaces in semi-dry conditions. Detected changes in metal tolerance were not associated with increased antibiotic tolerance indicating reduced risk of cross-selection or co-regulation of both tolerance mechanism in the exposure conditions. Cyclic exposure to copper and silver surfaces also did not increase mutation accumulation in the exposed populations indicating reduced risk of pathogenicity-associated hypermutator emergence on antimicrobial copper and silver surfaces compared to steel and glass in application-relevant semi-dry conditions.

Mutations in several known metal tolerance (*ompC*, *zraS*, *cysN*), stress response (*rssB, ybiB*) and cell wall integrity (*yfhM*) associated genes were detected in evolved populations with increased copper or silver tolerance. The abundance of mutations in sulfur and sulfur amino acid (*tnaA, ydjN, cysN*) or purine (*apt*) metabolism genes offering survival or growth benefit on copper and silver surfaces was surprising and should be studied further. Their effect could be plausibly explained by reduced metal uptake and increased metal detoxification by sulfidation or sequestering by adenine binding. Enriched mutations on copper and silver surfaces indicate a shift toward more energy-passive mechanisms of metal tolerance such as possible detoxication via sulfur metabolism (*tnaA, cysN*) or adenine accumulation (*apt*) and decreasing metal uptake (*ompC, ydjN*) in surface exposure. Although efflux-associated changes were not detected in the populations evolved on copper or silver surfaces, Sil-mediated efflux was rapidly activated by point mutations in *silS* gene after a single high-dose silver exposure during MBC assays indicating decreased risk of *Sil* activation in surface exposure conditions compared to ionic copper and silver in liquid medium. The *silS* mutants demonstrated substantial growth and survival benefit in MIC and MBC assays with ionic copper and silver as well as increased MIC of ampicillin and ciprofloxacin but did not provide a substantial protection against copper or silver surface exposure in semi-dry conditions. This suggests that efflux-based systems may help to maintain metal homeostasis in liquid medium but is of less relevance in environmentally exposed conditions. Furthermore, it could mean reduced risk of activation of the metal tolerance loci and possible co-selection of antibiotic resistance on antimicrobial metal surfaces in case of extrachromosomal genetic elements carrying both efflux-based metal tolerance loci and antibiotic resistance genes.

## Availability of data

Sequencing data is made available in European Nucleotide Archive (ENA), accession PRJEB75034. Raw numerical data is included figure by figure in Supplementary Data file. Any other inquiries can be directed to the corresponding author.

## Author contribution

MR and SP, experimental design. SP, SU and MR, lab experiments and analysis of the results. SP, MR and AI, interpretation of the results. MR writing of the manuscript. MR and AI editing of the manuscript. AI project management and funding acquisition.

## Supporting information

Supplementary Results file

Supplementary Raw Data file

