## Supplementary Figures and Tables for "Experimental evolution of *Escherichia coli* on semi-dry silver, copper, stainless steel, and glass surfaces"

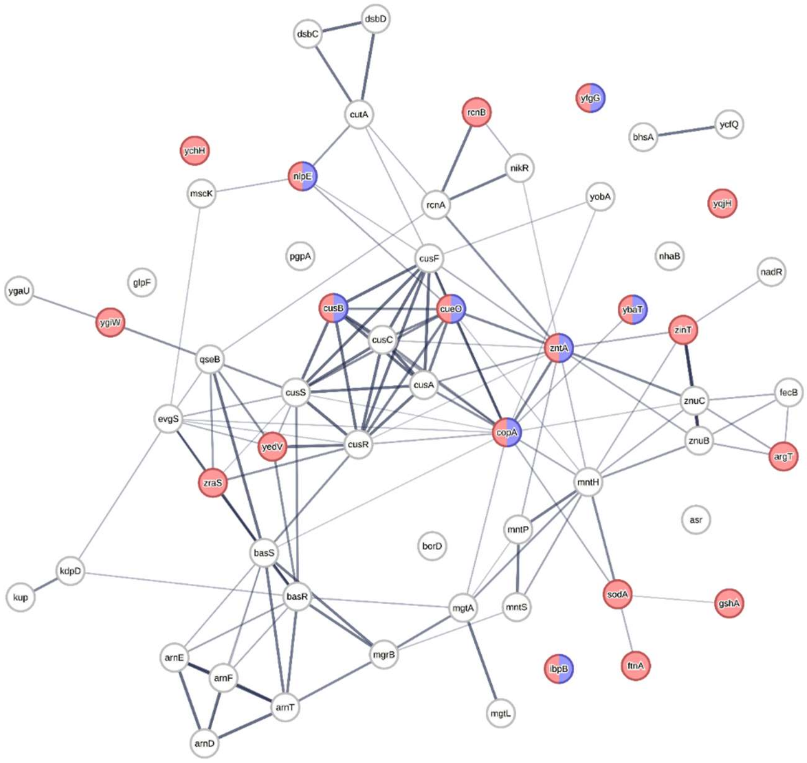


**Supplementary Figure S1.** Known and predicted interactions of the *E. coli* genes with relevant Gene Onthology (GO) functional annotations [136] retrieved from EMBL STRING [137], default settings. Interactions of the genes belonging to the GO term “response to metal ion” (GO:0010038). Overlapping genes with the GO term “response to stress” (GO:0006950) and “stress response to metal ion (GO:0097501) are marked in red and blue, respectively. Line thickness indicates the strength of data support for the interaction.

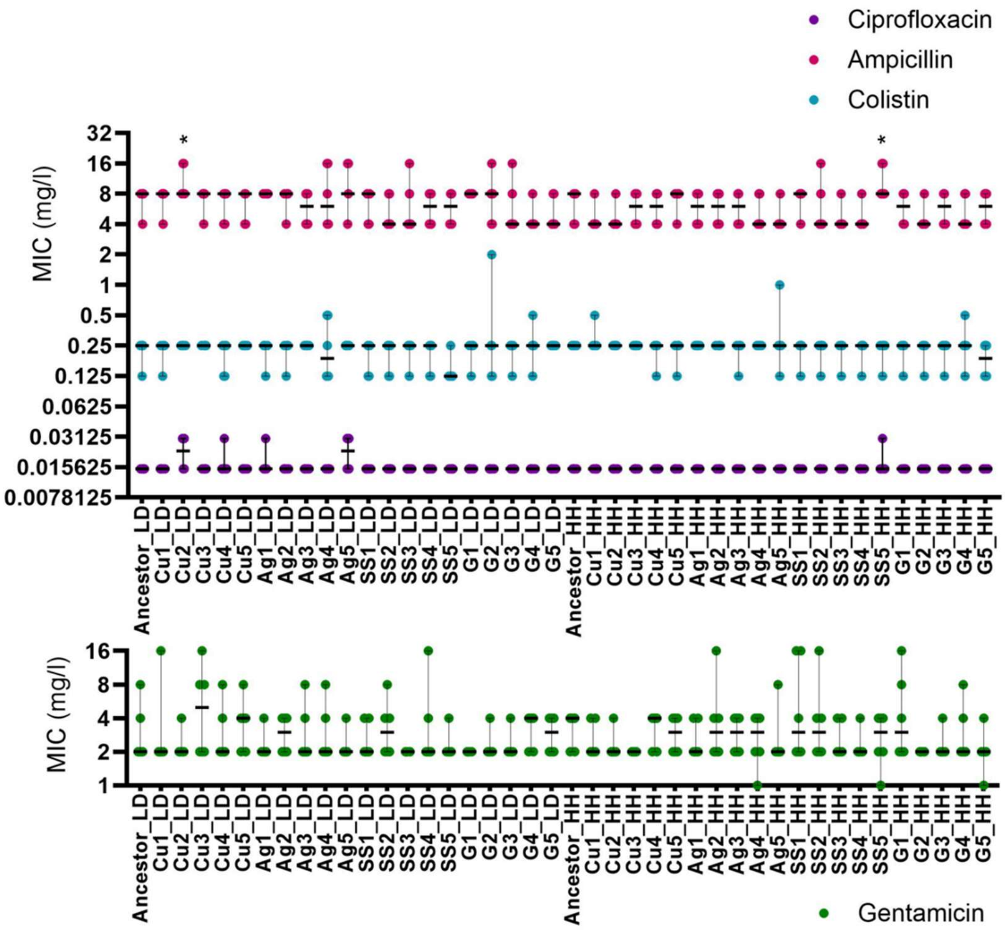


**Supplementary Figure S2. Antibiotic tolerance of the populations evolved in low-organic dry (LD) and high-organic humid (HH) and their respective ancestors, minimal inhibitory concentration (MIC).** Values for populations evolved on copper (Cu), silver (Ag), stainless steel (SS) and glass (G) are presented. Values of at least 6 data points are presented with median and range. Statistical significance of the difference of a single evolved lineage from the ancestor exposed in the same conditions is marked with * P< 0.05.

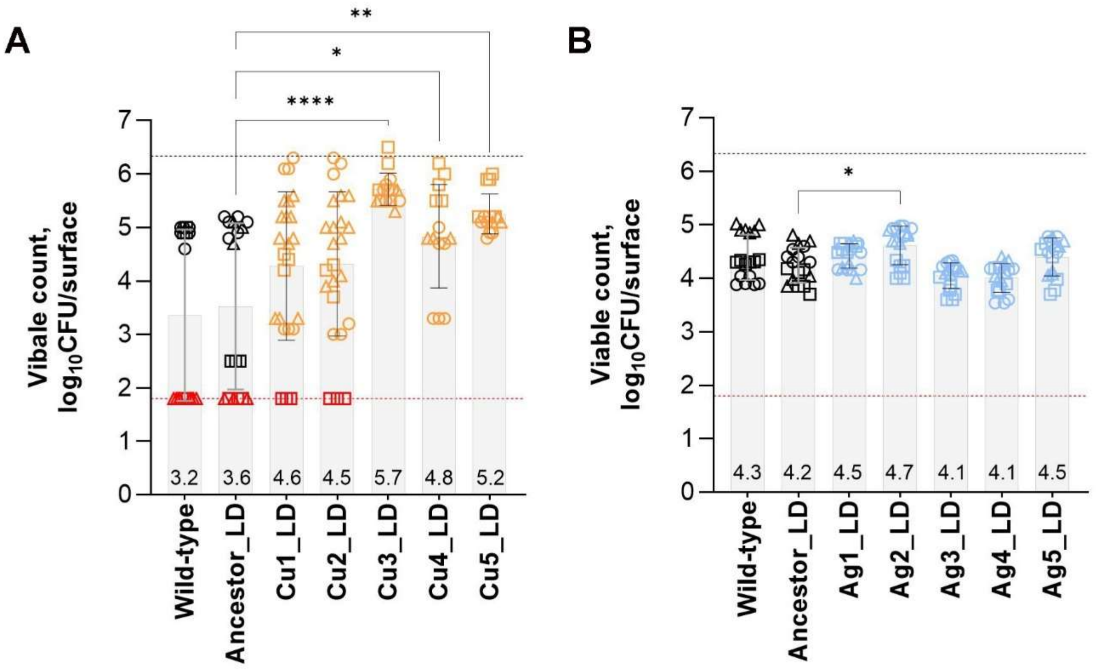


**Supplementary Figure S3. Copper (A) and silver (B) surface tolerance of the populations evolved in low-organic dry (LD) conditions.** Evolved populations were tested against the ancestral strain in the same conditions as in the corresponding evolution experiment. Values from three independent experiments are presented with mean ±SD. Median value is presented numerically. Symbol shapes indicate datapoints originating from different independent experiments. Limit of detection of viable counts is marked in red dotted line and red symbols indicate surfaces from which no colonies were retrieved. Black dotted line represents viable count in the initial inoculum. Statistical significance of the difference from the ancestor exposed in the same conditions is marked with * P< 0.05; ** P< 0.01; ****P< 0.0001 based on ANOVA followed by *post-hoc* testing for multiple comparisons.

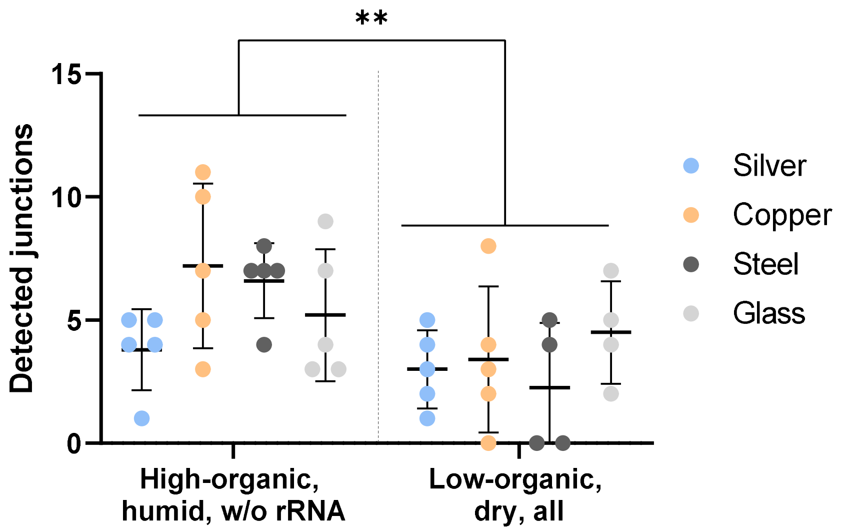


**Supplementary Figure S4. Total number of larger genomic rearrangements** (new junctions) detected in the WGS data of the evolved populations from high-organic humid and low-organic dry exposure conditions at 0.05 frequency after removing mutations present in the ancestor and rDNA reorganizations. ** P< 0.01

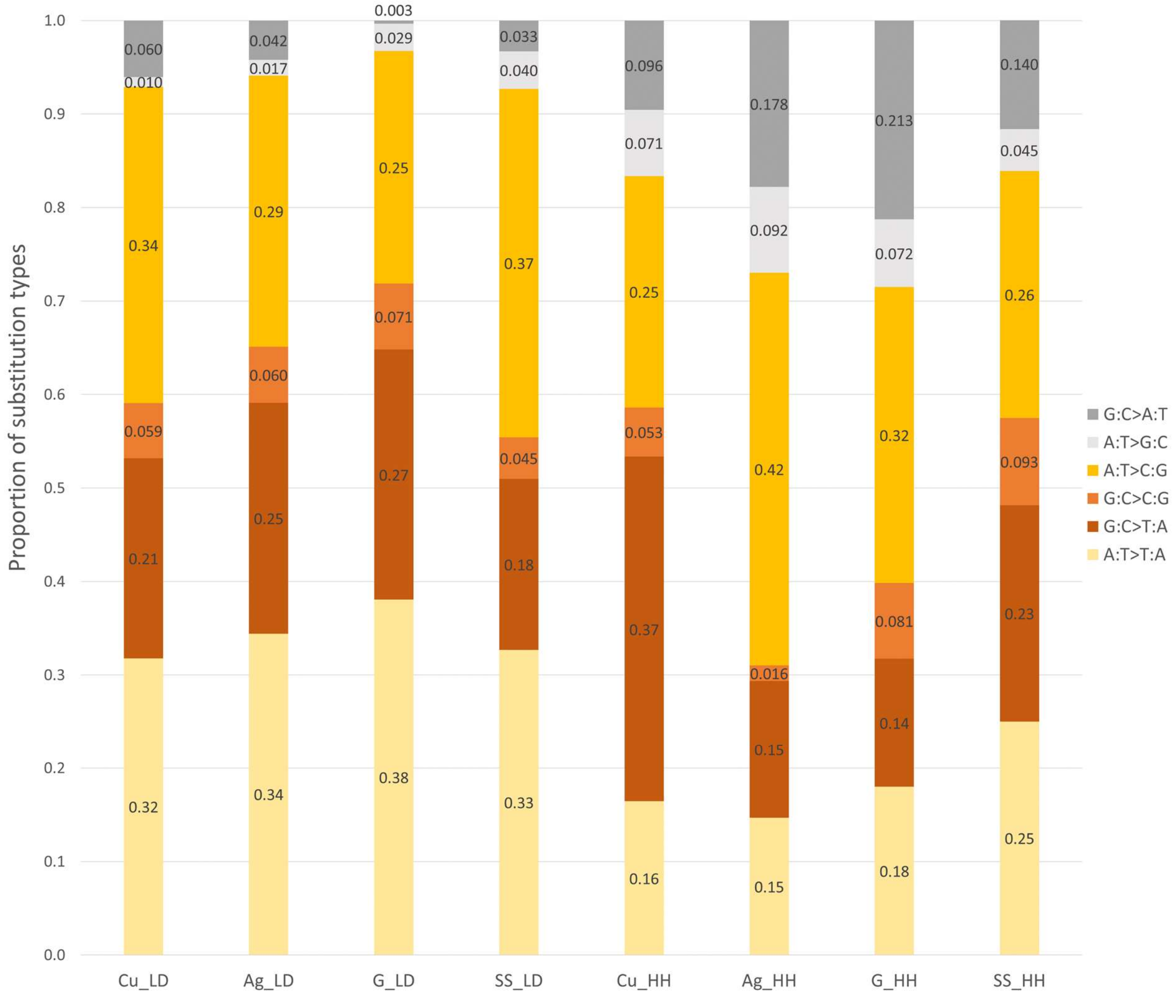


**Supplementary Figure S5.** Mean proportions of specific transitions and transversions in populations evolved on copper (Cu), silver (Ag), stainless steel (SS) and glass (G) in low-organic dry (LD) or high-organic humid (HH) exposure conditions.

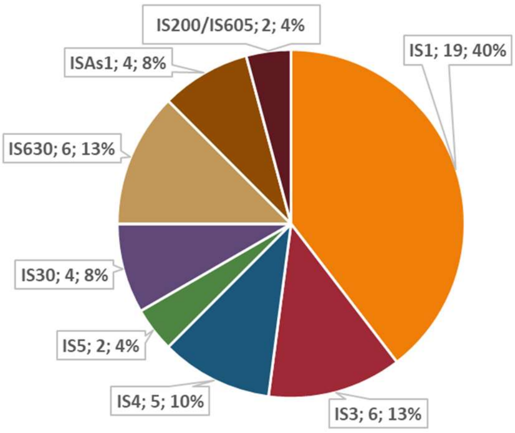


**Supplementary Figure S6**. 48 full length IS elements from 8 IS families were identified by ISfinder [44] in the reference genome of *E. coli* ATCC 8739 (GenBank: CP000946.1). Resequencing of the ancestral strain revealed a fixed mutation of 776 bp deletion confirming the loss of one full length IS1 (768 bp) from position 922 809 onwards. Most of the detected IS1 elements (14 out of 19) are most similar to IS1X2 with a deposited sequence originating from *Escherichia vulneris*.

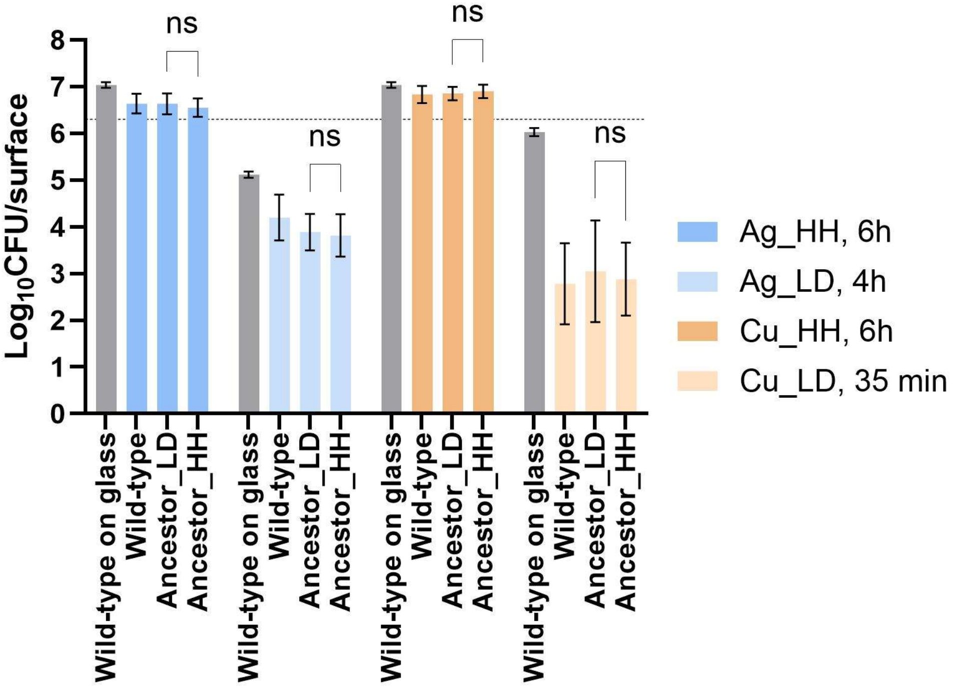


**Supplementary Figure S7. Viable counts of wild-type *E. coli* ATCC 8739 and ancestors of the low-organic dry (LD) and high-organic humid (HH) evolution experiments on silver (blue) and copper (orange) surfaces.** The four combinations of surface type (Ag or Cu) and exposure conditions (HH or LD) with the exposure time used for the combination are presented in the legend. Viability of wild-type *E. coli* on glass in the same conditions is presented in grey for comparison. No significant differences in viability (ns; P>0.05) of the ancestors was detected in any of the conditions tested (one-way ANOVA with post-hoc testing for selected multiple comparison, all comparisons shown on the figure).

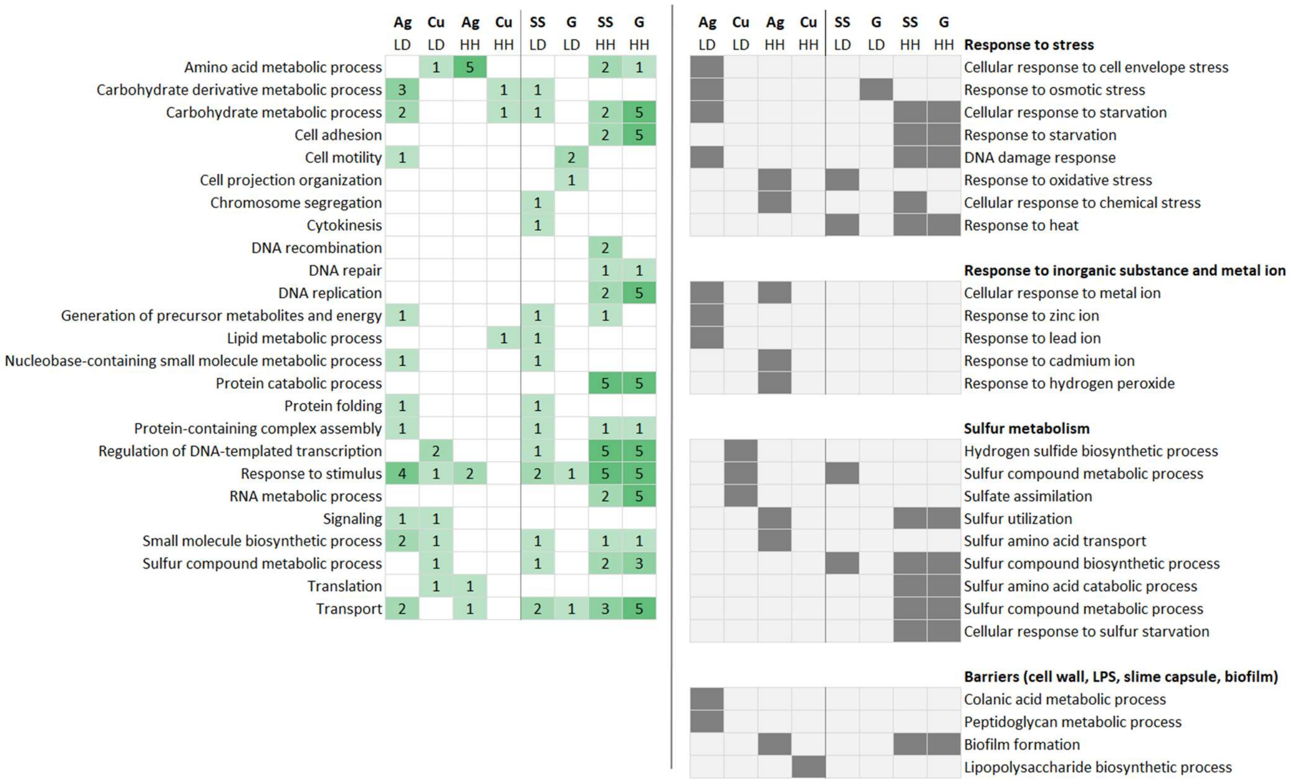


**Supplementary Figure S8. Functional annotation of genes potentially affected by mutations accumulated with ≥0.2 frequency** in population evolved on silver, copper, stainless steel (SS) and glass (G) in low-organic dry (LD) and high-organic humid (HH) exposure conditions. Number of parallel populations where mutations with shared GOslim_prokaryote annotations are presented in green gradient (left panel) and presence of more specific GO terms is presented as dark squares (right panel). QuickGO [138] was used to filter Gene Ontology database [136]. For copper and silver, occasionally occurring IS1-mediated deletions (*EcolC_2715*-…) also abundantly present on control surfaces were excluded from the analysis.

**A**

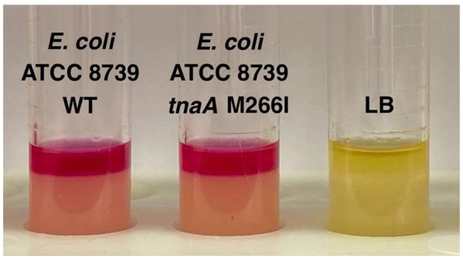


**B**

| | Soil load + cysteine<br>0, 54, 108, 216, 432, 864, 1728, 3456 mg Ag/l | Soil load + cysteine + ferric citrate<br>0, 54, 108, 216, 432, 864, 1728, 3456 mg Ag/l |
|---|---|---|
| WT,<br>2 h | 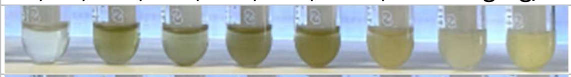 | 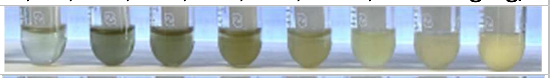 |
| *tnaA* M266I,<br>2 h | 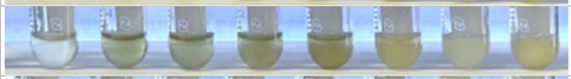 | 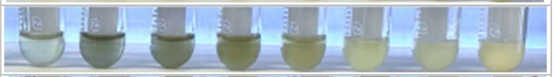 |
| WT,<br>24 h | 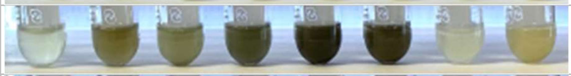 | 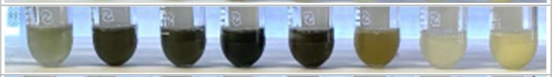 |
| *tnaA* M266I,<br>24 h | 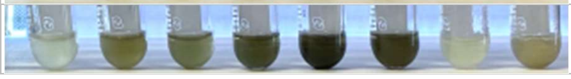 | 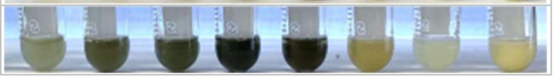 |

**Supplementary Figure S9**. Colorimetric detection of indole (A) and $H_2S$ (B)by wild-type *E. coli* ATCC8739 (WT) and its *tnaA* M266I mutant isolate. Both WT and its M266I mutant produce indole during overnight growth in LB (A) and $H_2S$ during 2 h and 24 h incubation in the presence and/or absence of $AgNO_3$ (B) confirming that the M266I mutation did not inactivate the tryptophanase enzyme. Quantitative colorimetric evaluation of $H_2S$ production (B) is complicated by the fact that not only the ferric salt, but also silver causes dark sulfide formation and higher turbidity caused by growth during Ag exposure at lower concentrations decreases perceived intensity of dark sulfides while Ag also causes growth inhibition at higher concentrations. However, a qualitative conclusion can be made that M266I mutation did not disrupt $H_2S$ production of *E. coli* with and without Ag presence.

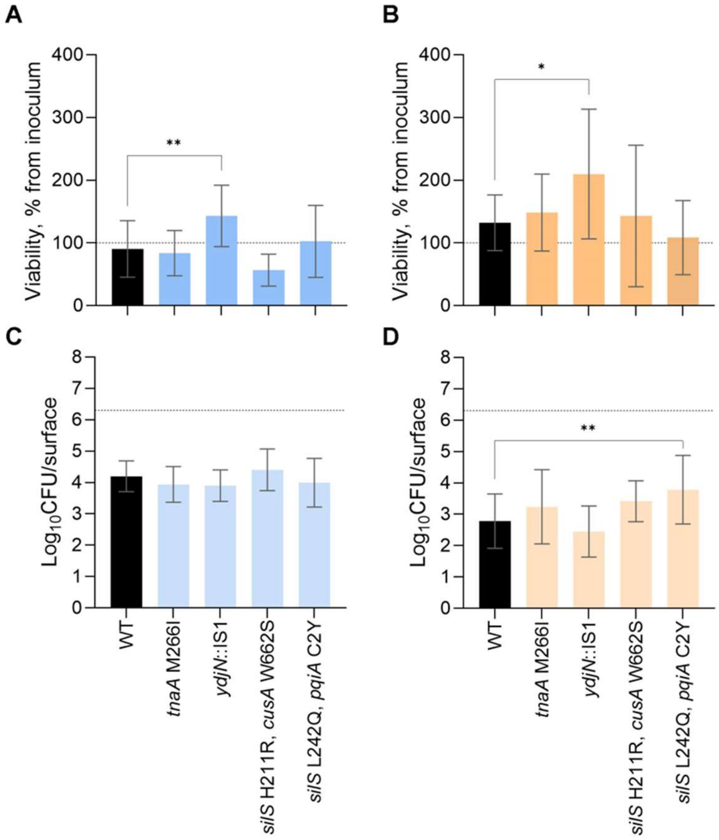


**Supplementary Figure S10. Silver (A,C) and copper (B, D) surface tolerance of the selected mutants and wild-type(WT) in high-organic humid (A,B) and low-organic dry (C,D) and conditions.** Isolates from Ag4_HH population carrying *ydjN*::IS1 or *tnaA* I266M mutations and *silS* mutants that emerged after the evolution experiment and were isolated from MBC assays of the evolved populations are presented. Values from three independent experiments (6 surface exposures per condition) are presented with mean ±SD. Ancestral viability (A,B) or pre-exposure inoculum baseline (C,D) is marked with a dotted line. Statistical significance of the difference from the ancestor exposed in the same conditions is marked with * $P < 0.05$; ** $P < 0.01$ based on non-parametric ANOVA followed by *post-hoc* testing for multiple comparisons.

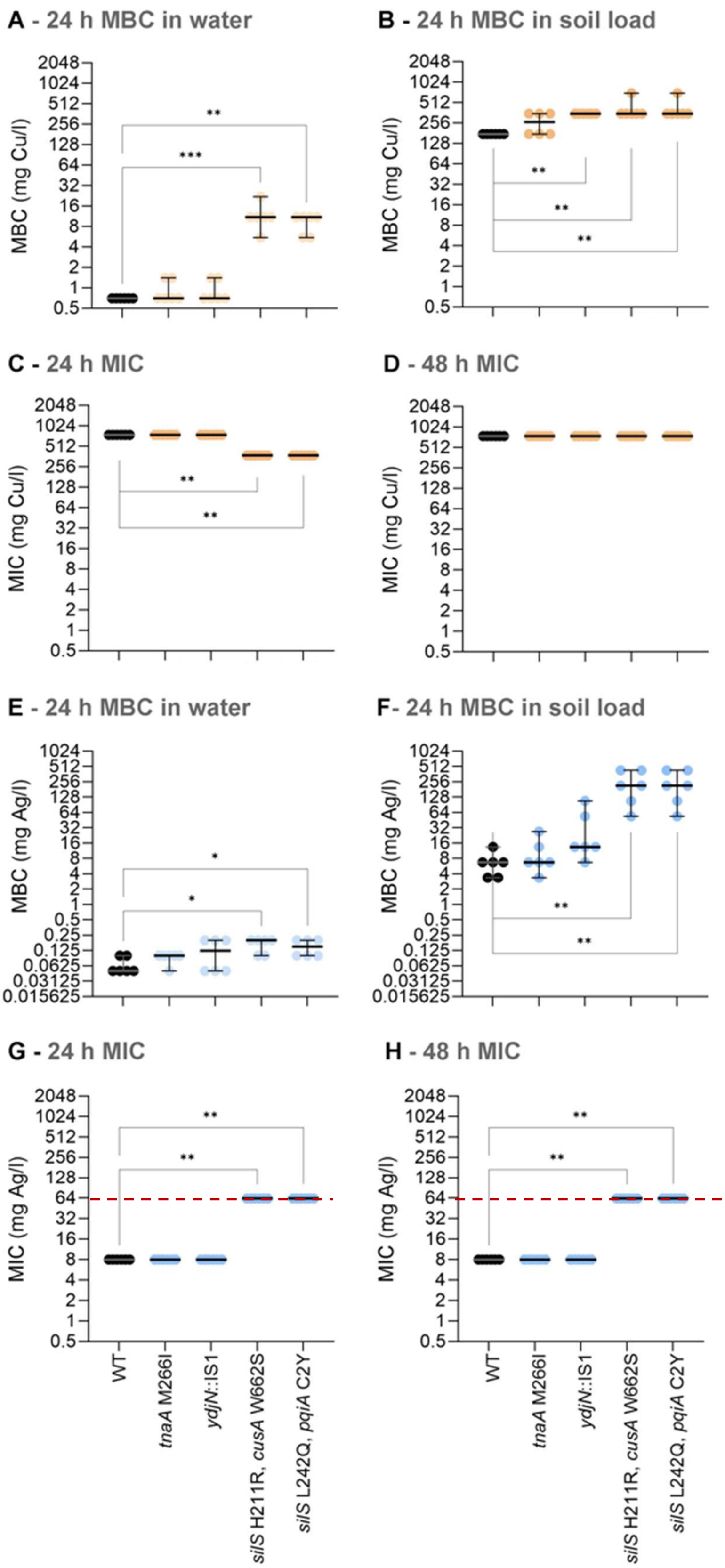


Highest Ag concentration tested for MIC, 64 mg/ml

**Supplementary Figure S11. MBC and MIC values of copper (A-D, orange) and silver (E-H, blue) salts of selected mutants and wild-type (WT).** MBC in water (A, E) and organic soil load (B, F) and MIC values at 24 h (C, G) and 48 h (D, H) are presented. On panels G and H MIC is ≥ 64 mg Ag/l as the detected value was the highest silver concentration tested in MIC assay. Isolates from Ag4_HH population carrying *ydjN*::IS1 or *tnaA* I266M mutations and *silS* mutants that emerged after the evolution experiment and were isolated from MBC assays of the evolved populations are presented. Values from three independent experiments with 6 data points are presented with median and range. Statistical significance of the difference from the ancestor exposed in the same conditions is marked with * P< 0.05; ** P< 0.01; *** P<0.001 based on non-parametric ANOVA followed by *post-hoc* testing for multiple comparisons.

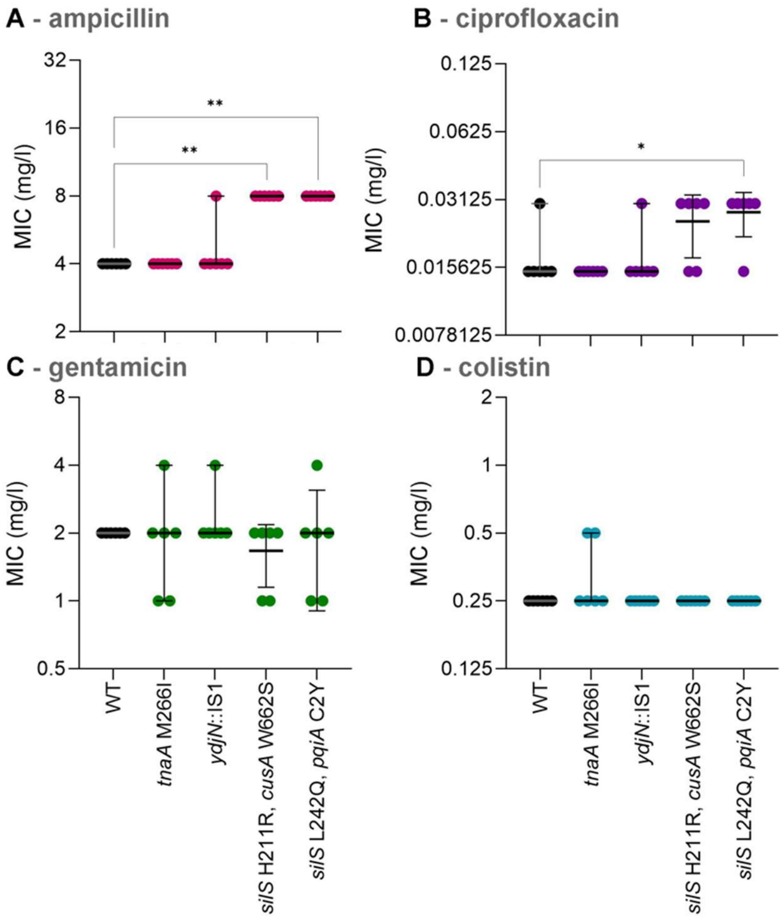


**Supplementary Figure S12. Ampicillin (A), ciprofloxacin (B), gentamicin (C) and colistin (D) tolerance of the selected mutants and wild type (WT).** Isolates from Ag4_HH population carrying *ydjN*::IS1 or *tnaA* I266M mutations and *silS* mutants that emerged after the evolution experiment and were isolated from MBC assays of the evolved populations are presented. Values from three independent experiments with 6 data points are presented with median and range. Statistical significance of the difference from wild-type exposed in the same conditions is marked with * P< 0.05; ** P< 0.01 based on non-parametric ANOVA followed by *post-hoc* testing for multiple comparisons.

**Supplementary Table S1**. Mutations accumulated in five parallel populations evolved on glass and steel surfaces in low-organic dry (LD), and high-organic humid (HH) exposure conditions. Presence of mutated genes also affected in silver and/or copper exposed populations is indicated for comparison. Frequency of the mutation in each population is shown numerically; only mutations at ≥0.2 frequency are presented. Grey text indicates mutation frequency <0.2 in cases where other mutations affecting the same gene also occur at ≥0.2 frequency. Synonymous SNVs, rRNA reorganizations and intergenic changes downstream of reading frames are excluded from the table. Full mutational profiles of all ancestors evolved populations and isolates can be found in Supplementary Results file.

| | | | | GO terms; Response to: | | Glass-exposed lineages | | | | | | | | | Steel-exposed lineages | | | | | | | | | Silver-exposed lineages | | | | | | | | | | Copper-exposed lineages | | | | | | | | | |
|---|---|---|---|---|---|---|---|---|---|---|---|---|---|---|---|---|---|---|---|---|---|---|---|---|---|---|---|---|---|---|---|---|---|---|---|---|---|---|---|---|---|---|---|
| | | | | | | G_LD | | | | G_HH | | | | | SS_LD | | | | SS_HH | | | | | Ag_LD | | | | | Ag_HH | | | | | Cu_LD | | | | | Cu_HH | | | | |
| Mutation | Annotation | Affected loci (changes upstream of and inside of protein-coding reading frames) | Gene (KEGG similarity) | Stress | Metal ions | 1 | 2 | 3 | 4 | 1 | 2 | 3 | 4 | 5 | 1 | 2 | 4 | 5 | 1 | 2 | 3 | 4 | 5 | 1 | 2 | 3 | 4 | 5 | 1 | 2 | 3 | 4 | 5 | 1 | 2 | 3 | 4 | 5 | 1 | 2 | 3 | 4 | 5 |
| T→C | intergenic (-824) | EcolC_0041 | *setC* | yes | no | | | | | | | | | | | | | | | 0.4 | | | | | | | | | | | | | | | | | | | | | | | |
| T→G | intergenic (-828) | EcolC_0041 | | | | | | | | | | | | | | | | | | 0.4 | | | | | | | | | | | | | | | | | | | | | | | |
| T→A | intergenic (-829) | EcolC_0041 | | | | | | | | | | | | | | | | | | 0.4 | | | | | | | | | | | | | | | | | | | | | | | |
| T→C | intergenic (-849) | EcolC_0041 | | | | | | | | | | | | | | | | | | 0.3 | | | | | | | | | | | | | | | | | | | | | | | |
| Δ38 bp | intergenic (-19 to -58/-114 to - | EcolC_0158/EcolC_0159 | *transposase/h* | -/no | -/no | | | | | | | | | | | | | | | | 1.0 | | 0.1 | | | | | | | | 0.1 | | 0.1 | | | | | | | | | | |
| C→A | A15S (GCA→TCA) | EcolC_0633 | *dnaG* | no | no | | | | | 0.3 | 0.1 | 0.1 | 0.1 | 0.6 | | | | | 0.3 | | | | 0.1 | | | | | | | | | | | | | | | | | | | | |
| A→C | F231V (TTT→GTT) | EcolC_0772 | *metK* | no | no | | | | | 0.2 | 0.2 | | | 0.2 | | | | 0.2 | | | | | 0.1 | 0.3 | 0.3 | 0.3 | | | 0.2 | | | | 0.2 | | | | | 0.2 | 0.2 | | 0.2 | | |
| A→T | E399D (GAA→GAT) | EcolC_0779 | *yggP* | no | no | | | | 0.6 | | | | | | | | | | | | | | | | | | | | | | | | | | | | | | | | | | |
| ::IS3 (+) | coding (457-459/1623 nt) | EcolC_0844 | *ygeP* | no | no | | | | | 0.6 | | | | | | | | | | | | | | | | | | | | | | | | | | | | | | | | | |
| C→A | A11S (GCC→TCC) | EcolC_1058 | *alpA* like | - | - | | | | | | | | | | | | | | | 0.3 | | | | | | | | | | | | | | | | | | | | | | | |
| T→A | intergenic (-105) | EcolC_1116 | *yfhH* | no | no | | | | | | | | | | | 0.3 | | | | | | | | | | | | | | | | | | | | | | | | | | | |
| T→A | intergenic (-104) | EcolC_1117 | *pgpC;yfhB* | no | no | | | | | | | | | | | 0.3 | | | | | | | | | | | | | | | | | | | | | | | | | | | |
| T→A | intergenic (-419) | EcolC_1509 | *yohG;mdtQ* | no | no | | | | 0.2 | | | | | | | | 0.2 | 0.2 | 0.1 | | | | | | | | 0.2 | | | | | | | | 0.2 | | | | | | | | |
| A→C | intergenic (-87) | EcolC_1720 | *dcyD* | yes | no | | | | | 0.1 | | | | | | | | | 0.6 | | | | 0.1 | | | | | | | | | | | | | | | | | | | | |
| ::IS4 (+) | coding (112-122/360 nt) | EcolC_1836 | *yeaR* | no | no | | | | | | | | | | | | | | | | 0.9 | | 0.1 | | | | | | | | 0.1 | | 0.1 | | | | | | | | | | |
| T→G | F61C (TTT→TGT) | EcolC_2105 | phage protein | - | - | | | 0.2 | 0.2 | | | 0.2 | | | | 0.2 | | | 0.2 | | | 0.2 | | | | | | | | | | | 0.2 | | | | | | 0.1 | | | | |
| T→G | E164A (GAA→GCA) | EcolC_2385 | *oppA* | yes | no | | | | | | | | | | | | | 0.2 | | | | | | | | | | | | | | | | | | | | | | | | | |
| Δ5770 bp | intergenic (+389) to coding (33 | EcolC_2386-EcolC_2389 | | | | | | | | | | | | | | 0.9 | | | | | | | | | | | | | | | | | | | | | | | | | | | |
| min Δ4,593 bp | IS1X2-mediated (EcolC_2454) | EcolC_2447-EcolC_2453 | | | | | | | | | | | | | | | 0.3 | | | | | | | | | | | | | | | | | | | | | | | | | | |
| min Δ2,116 bp | IS1X2-mediated (EcolC_2454) | EcolC_2455-EcolC_2456p | *ymgD* | no | no | | | | | | | | | | | | | | | 0.2 | | | | | | | | | | | | | | | | | | | | | | | |
| C→A | R73L (CGA→CTA) | EcolC_2521 | *flgH* | yes | no | 0.5 | | | | | | | | | | | | | | | | | | | | | | | | | | | | | | | | | | | | | |
| Δ1 bp | coding (97/660 nt) | EcolC_2528 | *flgA* | no | no | | | 0.2 | | | | | | | | | | | | | | | | | | | | | | | | | | | | | | | | | | | |
| Δ636 bp | coding (1842/2424) to coding ( | EcolC_2572-EcolC_2573 | *pgaA/pgaB* | no | no | | | | | 0.3 | 0.1 | 0.1 | 0.2 | 0.6 | | | | | 0.3 | | | | 0.1 | | | | | | | | | | | | | | | | | | | | |
| min Δ2,439 bp | IS1X2-mediated (EcolC_2714) | EcolC_2715-EcolC_2717 | *clpS, cspD* | yes,yes | no,no | | | | | | | | | | | | | | | | | | | | | | | | | | | | | 0.9 | | | | | | | | | |
| min Δ 3,459 bp | IS1X2-mediated (EcolC_2714) | EcolC_2715-EcolC_2718 | *clpS, cspD* | yes,yes | no,no | | | | | | | | | | | | | | | | | 0.3 | | | | | | | | | | | | | | | | | | | | | |
| min Δ5,750-7,386 bp | IS1X2-mediated (EcolC_2714) | EcolC_2715-EcolC_2720 | *clpS, cspD* | yes,yes | no,no | | | | | | | | | | | | | | 1.0 | 0.8 | | 0.8 | | | | | | | | | | 0.4 | | | | | | | | | 0.2 | | |
| min Δ8,880-9,231 bp | IS1X2-mediated (EcolC_2714) | EcolC_2715-EcolC_2722 | *clpS, cspD* | yes,yes | no,no | | | | | | 1.0 | 0.8 | 0.9 | | | | | | | | 1.0 | | | | | | | | | | | | | | | | | | | | | | |
| min Δ17,609 bp | IS1X2-mediated (EcolC_2714) | EcolC_2715-EcolC_2728 | *clpS, cspD* | yes,yes | no,no | | | | | 0.8 | | | | | | | | | | | | | | | | | | | | | | | | | | | | | | | | | |
| min Δ32,015 bp | IS1X2-mediated (EcolC_2714) | EcolC_2715-EcolC_2745 | *clpS, cspD* | yes,yes | no,no | | | | | | | | | | | | | | | | | | | | | | | | | | | | | | | | 1.0 | | | | | | |
| min Δ35,461 bp | IS1X2-mediated (EcolC_2714) | EcolC_2715-EcolC_2749 | *clpS, cspD* | yes,yes | no,no | | | | | 0.2 | 0.1 | | 0.1 | 0.6 | | | | | 0.3 | | | | 0.1 | | | | | | | | | | | | | | | | | | | | |
| min Δ13972 bp | IS1R-mediated (EcolC_2872) | EcolC_2857-EcolC_2870 | | | | | | | | | | | | | | | | | | | | | 0.2 | | | | | | | | | | | | | | | | | | | | |
| Δ9,624 bp | IS1R-mediated (EcolC_2872) | EcolC_2863-EcolC_2870 | | | | | | | | | 0.8 | | | | | | | | | | | | | | | | | | | | | | | | | | | | | | | | |
| G→A | A25T (GCC→ACC) | EcolC_2971 | *ybfE* | yes | no | | | | | | | | | | | | | | 0.6 | | | | | | | | | | | | | | | | | | | | | | | | |
| G→A | R312W (CGG→TGG) | EcolC_2996 | *djlC* | no | no | | | | | | | 0.7 | 0.6 | | | | | | | | | | | | | | | | | | | | | | | | | | | | | | |
| T→A | L242Q (CTG→CAG) | EcolC_3140 | *aes* | no | no | | | | | | | | | 0.1 | | | | | | 0.6 | | 0.1 | | | | | | | | | | 0.1 | 0.1 | | | | | | | | | | |
| ::IS1X2 (+) | coding (315-323/678 nt) | EcolC_3242 | *aroM* | no | no | | | | 0.6 | | | | | | | | | | | | | | | | | | | | | | | | | | | | | | | | | | |
| C→T | P340L (CCT→CTT) | EcolC_4070 | *gldA* | no | no | | | | | | | | | | | | | | | | | | 0.3 | | | | | | | | | | | | | | | | | | | | |

**Supplementary Table S2**. Differences in mutational profile of mutant *ydjN*, *tnaA* and *silS* isolates characterized in the study. Mutations present already in the evolved population of origin of each isolate are marked with blue background, mutations present in the Ancestor_HH, but not Ancestor_LD with green background and mutations emerged after the evolution experiments and not present in the evolved population of origin of each isolate on white background in the first 4 columns. Synonymous SNVs, rRNA reorganizations and intergenic changes downstream of neighboring reading frames are excluded from the table. Full mutational profiles of all ancestors, evolved populations and isolates can be found in Supplementary Results file.

| **Position** | **Annotation** | **Affected loci** | **Affected genes by similarity** | ***tnaA* M266I** | ***ydjN* ::IS1** | ***silS/pqiA*** | ***silS/cusA*** |
|---|---|---|---|---|---|---|---|
| 1,900,542 | W8* (TGG→TGA) | EcolC_1701 | *fliI* | + | + | + | |
| 2,624,634 | Δ3 bp (855-857/1677 nt) | EcolC_2385 | *oppA* | + | + | + | |
| 2,625,746 | intergenic A→C (-256) | | | + | + | + | |
| 2,987,583 | Δ7339 bp | EcolC_2715-EcolC_2721 | *clpS,cspD,...* | + | | | |
| **4,740,468** | **M266I (ATG→ATT)** | **EcolC_4286** | ***tnaA*** | **+** | | | |
| 3,098,445 | A17V (GCG→GTG) | EcolC_2832 | *glnH* | | + | | |
| **2,105,539** | **::IS1 (209-217/1392 nt)** | **EcolC_1903** | ***ydjN (tcyP)*** | | **+** | | |
| 1,230,172 | intergenic G→A (-112) | EcolC_1127 | *yphH* | | | + | |
| | intergenic G→A (-6) | EcolC_1128 | *yphG* | | | + | |
| **2,898,355** | **C2Y (TGC→TAC)** | **EcolC_2646** | ***pqiA*** | | | **+** | |
| **3,746,708** | **L242Q (CTG→CAG)** | **EcolC_3427** | ***silS*** | | | **+** | |
| 4,548,981 | S102C (AGC→TGC) | EcolC_4122 | *fdhD* | | | + | |
| 1,755,361 | intergenic T→A (-47) | EcolC_1578 | *yegH* | | | | + |
| | intergenic T→A (-612) | EcolC_1579 | *wza* | | | | + |
| **3,356,525** | **W662S (TGG→TCG)** | **EcolC_3071** | ***cusA*** | | | | **+** |
| **3,746,615** | **H211R (CAC→CGC)** | **EcolC_3427** | ***silS*** | | | | **+** |
